# Biofilm-deficient mutants of *Pseudomonas aeruginosa* have wild-type levels of antibiotic tolerance in a model of cystic fibrosis lung infection

**DOI:** 10.64898/2026.02.17.706333

**Authors:** Jenny L Littler, Niamh E Harrington, Dean Walsh, Ramón Garcia Maset, Saskia E. Bakker, Christopher Parmenter, Freya Harrison

**Affiliations:** School of Life Sciences, University of Warwick, Coventry CV4 7AL UK; Institute of Infection, Veterinary and Ecological Sciences, University of Liverpool, L69 3BX, UK; Blizard Institute, The Faculty of Medicine and Dentistry, Queen Mary University of London, London E1 2AT, UK; Warwick Medical School, University of Warwick, Coventry CV4 7AL UK; Institute of Biomedical Engineering, Department of Engineering Science, University of Oxford, Oxford, UK; Advanced Bioimaging Research Technology Platform, University of Warwick, Coventry CV4 7AL UK; The Nanoscale and Microscale Research Centre, University Park Campus, University of Nottingham, NG7 2RD, UK

## Abstract

Opportunistic, biofilm-forming pathogens such as *Pseudomonas aeruginosa* can employ an array of strategies to reduce the impact of antibiotics on their survival. The biofilm matrix can prevent antibiotics from reaching bacteria embedded within it; general changes in metabolic activity alter susceptibility to specific drugs dependent on the target; changes in the membrane and the expression of channel or pump proteins embedded within it affect drug uptake and efflux; and production of antibiotic-degrading enzymes can remove the threat. In this study, we report that biofilm-deficient mutants of two well-studied lab strains of *P. aeruginosa* (PA14 and PAO1) have wild-type (WT) levels of tolerance to colistin and meropenem when allowed to establish mature populations in an *ex vivo* pig lung model of cystic fibrosis lung infection. This result was unexpected, given previous work in our lab that suggested physical protection of cells from colistin by the biofilm matrix was important for tolerance. The biofilm defects in the mutants were confirmed using electron and light microscopy, and cryo scanning electron microscopy was used to visualise the hydrated biofilm matrix in the WT. Using RNA sequencing of the PA14 WT and an isogenic mutant lacking the pel polysaccharide, we were able to identify a small number of differences in the responses of the two genotypes to the lung environment and to exposure to sub-bactericidal colistin in the lung model. Notably, there was differential upregulation of the MexXY-OprM and MexEF-OprN multidrug efflux pumps. Hence, mature biofilm structure is not essential for tolerance, as other mechanisms may be sufficient in this environment. However, the relative roles of biofilm matrix *versus* cellular changes in physiology in conferring antibiotic tolerance in this environment remain to be fully elucidated.

## 2 Introduction

Biofilms are aggregates of microbial cells, encased in a self-produced polymeric extracellular matrix that protects from environmental stressors; these aggregates may be attached to surfaces, or form as non-surface-attached aggregates in host fluids and tissues (Davies, 2002, Sauer et al., 2022). The bacterium *Pseudomonas aeruginosa* is a master biofilm former, and a major pathogen of the lungs of people with cystic fibrosis (CF) (Cystic Fibrosis Trust, 2025; Ledger et al. 2024). Its colonisation of CF lungs is associated with increased morbidity and mortality (Moreau-Marquis et al., 2008). Despite frequent use of antibiotics, these infections are difficult to eradicate, caused in part by the biofilm extracellular matrix acting as a barrier to treatment: some antibiotics cannot diffuse through the matrix, whilst others bind to components of the matrix (Olivares et al., 2020). Further, growth as a biofilm, and cues from the unique environment encountered within the CF lungs, drive changes in cellular metabolism and gene expression that increase antibiotic tolerance (Van den Bossche et al., 2021). Models that replicate this environment and therefore *in vivo*-like bacterial biofilm growth and physiology, allow us to understand how the environment alters the resistance profiles of bacteria and better understand pathogenicity and persistence (Grassi and Crabbe, 2024).

We previously showed a high level of tolerance to antibiotics by the *P. aeruginosa* laboratory strain PA14 when grown in an *ex vivo* model of CF lung biofilm, which comprises a section of pig bronchiole and synthetic CF sputum medium (*ex vivo* pig lung model, EVPL) (Harrington et al., 2020). In the case of the antibiotic colistin, we attributed this high tolerance to an inability of the drug to penetrate the biofilm matrix (Sweeney et al., 2021). However, changes in other aspects of cellular physiology, cued by the CF lung environment, may also lead to increased tolerance (Harrington et al., 2022, Huang et al., 2015, Palmer et al., 2007). We have since shown that EVPL biofilms have higher expression of two loci involved in resistance to cationic antimicrobial peptides, *arnA* and *crpP,* than planktonic cultures in synthetic CF mucus medium, which could contribute to heightened colistin tolerance (Harrington et al., 2022). In addition, other researchers have found that while overproduction of the biofilm matrix can increase antibiotic tolerance, loss of matrix production does not necessarily lead to reduced tolerance if bacterial motility is restricted (Goltermann and Tolker-Nielsen, 2017, Staudinger et al., 2014). Therefore, we set out to explore the role of a key biofilm matrix polymer in *P. aeruginosa* antibiotic tolerance in the EVPL model, by comparing the tolerance of PA14 wild-type (WT) bacteria and the respective biofilm-deficient mutants.

Exopolysaccharides are a key component of the extracellular matrix of biofilms. *P. aeruginosa* strains can produce up to three: pel, psl and alginate. The pel and psl polysaccharides are important for the early development of biofilms, giving the matrix its structural support, as well as maintaining the balance of water and nutrients (Zeng et al., 2015). Alginate is only produced by some strains of *P. aeruginosa*, causing a mucoid phenotype and contributing to the protection of the cell and surface adhesion (Wozniak et al., 2003b). Clinical *P. aeruginosa* isolates from CF patients commonly show increased production of pel, psl and alginate as infection progresses (Harrison et al., 2020). Overproduction of these exopolysaccharides in a lab strain via deletion of *mucA* (leading to overproduction of alginate and mucoidy), *wspF* or *yfiR* (both leading to overproduction of pel and psl and a rugose small-colony phenotype) has been linked to increased tobramycin and ciprofloxacin resistance. These phenotypescan be reversed to normal colony morphology and WT or near-WT levels of antibiotic sensitivity by deleting the loci essential for biosynthesis of alginate in the Δ*mucA* background, or pel and psl in the Δ *wspF* or Δ*yfiR* background. (Goltermann and Tolker-Nielsen, 2017). However, the same study also showed that WT lab isolates and isogenic mutants lacking exopolysaccharide biosynthesis could have comparable levels of antibiotic tolerance if bacterial aggregation was enforced by growth in an agar gel. Therefore, it is not clear how much protection the exopolysaccharide matrix gives to the bacterial population against antimicrobials. This is a particularly pertinent question in the CF lung environment, which can directly reduce antibiotic efficacy (e.g. via mucin binding or reduced pH (Van den Bossche et al., 2021)). Additionally, growth in media supplemented with CF sputum has been shown to cue expression of bacterial efflux pumps and antibiotic- degrading enzymes (Martin Lois et al., 2018, Drevinek et al., 2008), and growth in synthetic CF sputum medium and our EVPL model has been shown to lead to increased expression of antimicrobial resistance pathways (Harrington et al., 2022).

To understand more clearly the role that the extracellular matrix plays in biofilm resistance, we investigated the growth and antibiotic tolerance of *P. aeruginosa* mutants that are compromised in biofilm matrix polysaccharide production in the EVPL model of CF lung biofilm. We have previously demonstrated that growth of a *pelA^−^* transposon mutant of the commonly studied lab strain PA14 is comparable to the WT in terms of viable cell numbers, but the mutant has reduced biofilm matrix formation that lacks a mature structure in the EVPL model; i.e. it forms a bacterial population in the model but not a true biofilm (Harrington et al., 2020). PA14 lacks the genes for psl production and thus relies solely on pel for biofilm structural development. The production of pel is coordinated by the *pelABCDEFG* operon (Ghafoor et al., 2013). The periplasmically localised protein PelA deacylates the pel polymer whilst in the periplasm, a required step for maturation and secretion (Colvin et al., 2013). Loss of the *pelA* gene has been shown to significantly reduce the ability of PA14 to produce a mature biofilm on solid surfaces (Friedman and Kolter, 2004).

In the present work, we compared the susceptibility of EVPL-grown populations of WT and the *pelA^−^* mutant to clinically relevant antibiotics. Surprisingly, we found that the mutant populations were just as tolerant to colistin and meropenem as the WT biofilm populations. Comparable results were found when we compared the pel- and psl- producing lab strain PAO1 with a *pelApslBCD* deletion mutant, showing this effect is not strain specific. We therefore concluded that, whilst the biofilm matrix and structure can significantly increase antibiotic tolerance, this may not be the only fundamental factor in high-level antibiotic tolerance/resistance in the lung microenvironment. To begin to explore the mechanistic basis for these results, we then focussed on the PA14 WT/mutant pair and the antibiotic colistin. We used RNA-seq to study the impact of reduced pel production on the transcriptome of *P. aeruginosa* PA14 when grown in the EVPL model vs in synthetic CF sputum medium alone, and in the EVPL model with and without exposure to sub-bactericidal colistin. Our results suggest that in this environment, mechanisms such as close bacterial packing and efflux pump expression may suffice to confer antibiotic tolerance in the absence of mature matrix. This observation can be built on to dissect the relative roles of biofilm matrix and non-matrix-dependent tolerance mechanisms in the CF lung in future.

## 3 Materials and methodology

### 3.1 Bacterial isolates and growth conditions

*P. aeruginosa* PA14 wild type (WT) was compared with an isogenic insertional mutant containing the MAR2xT7 transposon in *pelA* (mutant ID 26187 from the PA14 Non-Redundant Transposon Insertion Mutant Set (Liberati et al., 2006)). The location of the transposon in the *pelA* locus was confirmed by PCR and Sanger sequencing following the instructions in the user manual for the mutant library (available at https://pa14.mgh.harvard.edu/cgi-bin/pa14/downloads.cgi). This mutant is hence referred to as PA14 *pelA^−^*. PAO1 WT and an isogenic *pelApslBCD* deletion mutant were kindly supplied by Matthew Parsek (Borlee et al., 2010). This mutant is henceforth referred to as PAO1 *△pelpsl.* Bacteria were cultured on Luria-Bertani (LB) agar plates (LB broth: Melford Laboratories; agar: Formedium), grown in static conditions at 37 ^◦^C. All culture steps below were conducted at 37 °C, without shaking.

The synthetic cystic fibrosis media (SCFM) (Palmer et al., 2005) was designed to replicate the sputum found in a cystic fibrosis lung. It includes a number of minerals, ions and amino acids. The recipe prepared was based on the published protocol by Palmer *et al* (Palmer et al., 2005) and adapted by removing glucose (Harrington et al., 2020). The standard lab media Mueller-Hinton broth (MHB) and cation-adjusted Mueller-Hinton broth (caMHB) (both Sigma-Aldrich) were prepared following the manufacturers’ instructions.

### 3.2 The *ex vivo* pig lung model

The *ex vivo* pig lung model (EVPL) was developed to replicate the lungs environment in cystic fibrosis (CF). Pig lungs were obtained from a local butcher (S Quigley butchers, Cubbington or Taylors, Coventry) and the bronchiolar tissue was dissected following a procedure established by Harrison and Diggle (2016) (Harrison et al., 2014). (An open-access video protocol is also available (Harrington and Harrison, 2023)) Briefly, the pig lungs were dissected using a sterile razor blade to make a vertical cut into the lung tissue to expose the bronchiolar tissue and cut away the lung tissue from the cartilage. This bronchiolar tissue was then removed, and any excess tissue was removed from the cartilage. The bronchiolar tissue was then placed into a 50 ml falcon tube containing a 1:1 ratio of Roswell Park Memorial Institute (RPMI) 1640 medium and Dulbecco’s modified Eagle medium (DMEM) (both from Sigma Aldrich) supplemented with 50 *µ*g ml^-1^ ampicillin (Sigma-Aldrich), this was left for 10 minutes at room temperature. Bronchiolar cartilage was then cut into 5mm longitudinal strips and these were again washed in fresh 1:1 mix of RPMI, DMEM and ampicillin. Finally, these strips were cut into 5mm x 5mm squares before being washed in RPMI and DMEM and finally in SCFM. These square pieces were then sterilised with shortwave ultraviolet (UV) light for 5 minutes (Carlton germicidal cabinet).

Tissue pieces were then transferred to a well of a flat bottomed 24-well plate (Corning Costar) containing a 400 *µ*l SCFM agar pad solidified with 0.8% agarose. A sterile 29 G hypodermic needle (Becton Dickinson Medical) was used to touch the surface of a colony from the LB plate it was grown on and then pierce a piece of bronchiolar tissue. Uninfected lung pieces were used as a negative control; these were mock inoculated with a sterile hypodermic needle. To further mimic the lung environment, 500 *µ*l of SCFM was added to each tissue well. The plate was covered with a sterilised Breathe-EASIER membrane (Diversified Biotech, USA). The plates were then incubated for 48 hours. Two lungs from different pigs were used as biological repeats per experiment, with at least three technical tissue replicates per condition.

To conduct antibiotic susceptibility testing in the EVPL, infected bronchiolar tissue pieces were incubated for 48 hours as above, then transferred to a 48 well plate containing 500 *µ*l of antibiotic diluted at the desired concentration in SCFM. The tissue pieces were incubated at 37 °C for a further 24 hours. Following incubation with the antibiotic, the tissue was removed from the plates with sterile forceps and washed in 500 *µ*l of PBS to remove non-biofilm associated cells. The lung tissue pieces were transferred to sterile homogenisation tubes containing eighteen 2.38 mm metal beads (Fisherbrand, UK) and 1 ml of PBS. The bacteria were recovered from the tissue-associated populations in the homogenisation tubes using a FastPrep-24 5G (MP Biomedicals) for 40 s at 4 m s. The homogenate was then serially diluted in PBS and plated on LB agar to determine bacterial load. Plates were incubated and colony forming units (CFU) per lung piece calculated from the colony counts.

### 3.3 Microscopy

#### 3.3.1 Light microscopy

EVPL infected with *P. aeruginosa* PA14 WT or *pelA^−^* was grown for 48 hours to allow the WT to establish a mature biofilm. Sections were fixed in 10% neutral buffered formalin and processed for wax embedding. 5 µm sections were mounted on slides and stained with haematoxylin-eosin, following our standard protocol (Harrington et al., 2020). Images were taken using a Zeiss Axio Scope.A1 light microscope with the Zeiss AxioCam ERc 5s and Zeiss ZEN Blue software.

#### 3.3.2 Scanning electron microscopy (SEM)

EVPL infected with *P. aeruginosa* PA14 WT or *pelA^−^*was grown for 48 hours to allow the WT to establish a mature biofilm. This was then washed with PBS and left in formaldehyde 4% aqueous solution (VWR) overnight to fix the sample. After the fixing step, 1 ml of PBS was used to wash the sample and remove any formaldehyde, this step was repeated 3 times. The sample was then dehydrated using increasing concentrations of ethanol. Ethanol was left on the sample for 1 hour before being removed and added to a higher concentration of ethanol. Ethanol concentrations used were 20%, 50%, 70%, 90%, 100% and 100%. Samples were then left in 0.5 mL of hexamethyldisilazane (electronic grade, 99+%; Thermo Scientific Alfa Aesar) for 1 hour, the remaining hexamethyldisilazane was removed and the sample was left to dry for 30 minutes in a laminar flow cabinet. The tissue was then coated with carbon twice. Images were obtained using a Zeiss Gemini electron microscope with an InLens detector, at 1kV. Omero version 5.7.1 was used to crop the microscopy images, add labels and add a scale bar.

#### 3.3.3 Cryo scanning electron microscopy (CryoSEM)

EVPL infected with *P. aeruginosa* PA14 WT or *pelA^−^*was grown for 48 hours to allow the WT to establish a mature biofilm. The samples were then washed in 1 mL of PBS and then fixed with 10% neutral buffered formalin (VWR Chemicals, UK). The tissues were then plunged into liquid nitrogen slush at cryogenic temperatures of -200 °C. Cryo-EM was performed by Saskia Bakker (University of Warwick, UK) and Chris Parmenter (University of Nottingham, UK). Cryo-SEM was performed at the Nanoscale and Microscale Research Centre (nmRC), using the Zeiss Crossbeam 550 SEM equipped with a Quorum 3010 Cryo-system. The samples were coated in platinum for 45-60 seconds at mA. Imaging was performed at 2 kV, with a current of 300pA at a working distance of 5 mm. Omero version 5.7.1 was used to crop the microscopy images, add labels and add a scale bar.

### 3.4 Statistical analysis of bacterial count data

All statistical analysis was performed in RStudio (Macintosh; Intel Mac OS X version 1.1.456) (RStudio, https://www.rstudio.com/) (R Core Team, 2021). Linear models were fitted and model assumptions tested by Q-Q plots and residuals vs. fits plot. An ANOVA was used to test general significance of independent variables, and Dunnett’s of Dunn’s test was used to compare treated samples vs the control. A Tukey post-hoc test was used when all samples were compared against each other. When a non-parametric test was needed, Mann-Whitney and Wilcoxon’s tests were used to assess significance, with a Bonferonni correction. P values <0.05 were considered significant.

### 3.5 RNA sequencing

#### 3.5.1 RNA extraction samples and conditions

PA14 WT and *pelA^−^* grown for two days in either the EVPL model or *in vitro* SCFM. *In vitro* samples consisted of *P. aeruginosa* strains PA14 WT and *pelA^−^* grown in 1 mL SCFM for 48 hours. After 48 hours of growth, either 64 *µ*g/mL of colistin or an equivalent volume of SCFM only was added to the EVPL model for 2 hours. Two lungs and 9 technical replicates per lung were used, 6 for RNA extraction and 3 for CFU counts. After 48 hours of growth, *in vitro* samples were added to fresh SCFM, without colistin, for 2 hours.

#### 3.5.2 RNA extraction

RNA was extracted from the samples described above using the phenol:chloroform method. Lung homogenate or SCFM culture was transferred to LoBind 2 ml tubes (Eppendorf) and one volume of killing buffer (20 mM Tris-HCl pH 7.5, 5 mM MgCl2, 20 mM NaN3) was added to each sample, these were centrifuged at 13,000rpm for one minute to pellet the cells, snap frozen in dry ice and kept at −80 ^◦^C for at least one hour. Samples were then defrosted on ice and the killing buffer removed. The pellets were resuspended in 600 *µ*l of LETS buffer (0.1 M LiCl, 0.01 M Tris-HCl pH 7.5, 0.2%), transferred into a lysing matrix B tube (MP Biomedicals, USA) and bead beaten using a FastPrep-24 5G (MP Biomedicals) for 3 cycles: 6.0m/s for 40 seconds followed by a 5-minute pause on ice. The tubes were then centrifuged for 10 minutes at 13,000 rpm. One volume of phenol:chloroform:isoamyl alcohol (PCI) (125:24:1) (Invitrogen) was added to each sample, vortexed at max speed ∼ 14,000 rpm for 5 minutes and then centrifuged at 15,000 rpm, 4 ^◦^C for 5 minutes. The top layer was then transferred to a new sterile RNase-free LoBind 2 mL tube (Sarstedt Ltd) and the phenol:chloroform:isoamyl steps were repeated as described above. Following this, the top layer was transferred to a new lo bind RNase-free 2 mL tube and 1 volume of chloroform:isoamyl alcohol 24:1 (Sigma-Aldrich) was added to each sample, vortexed at max speed for 5 minutes and centrifuged at 15,000 rpm at room temperature for 5 minutes. The top layer was again transferred to a new LoBind 2 mL tube, 1 volume of 100 % isopropanol (Fisher Scientific / Acros Organics) and 0.1 volume of sodium acetate (Fisher Scientific) were added to each sample. The samples were then inverted 6 times to mix and left at −20 ^◦^C overnight to precipitate the RNA.

The samples were then centrifuged at 15,000rpm at room temperature for 15 minutes. The supernatant was discarded and the pellet re-suspended in one volume of 70 % ethanol. This was then centrifuged again at 15,000 rpm at 4 °C for 15 minutes. The supernatant was removed and the pellets re-suspended in 50 µL RNA free water. 3 µL aliquot was taken for RNA concentration, the rest of the samples were snap frozen in alcohol plus dry ice and stored at −80 ^◦^C.

#### 3.5.3 Determining RNA concentration

Each sample had a 3 µL aliquot taken for RNA quantification using a Qubit RNA broad range (BR) Assay kit (Invitrogen) and Qubit 1.0 fluorometer (Invitrogen).

#### 3.5.4 RNA sequencing (outsourced)

Extracted RNA was sent to Source Bioscience, UK for sequencing. They performed gDNA removal, initial QC assessment, library preparation, rRNA removal and RNA sequencing. The samples were treated for rRNA depletion using NEBNext rRNA Depletion Kit (Bacteria) with NEBNext Ultra™ II Directional RNA Library Prep Kit for Illumina according to the manufacturer’s protocol. The initial RNA input was 150 ng. During this process, the libraries were indexed using NEBNext Multiplex Oligos for Illumina (96 Unique Dual Index Primer Pairs Set 4). Source Bioscience sequenced the data on an Illumina NovaSeq platform.

#### 3.5.5 RNA sequencing sample processing

RNA data processing was performed on the cloud infrastructure for big data microbial bioinformatics (CLIMB) server (Connor et al., 2016). The fastq files were initially quality checked using fastqc v0.11.8. Quality checks included GC content, read repeats, adapter contamination and sequence length. The QC step was repeated after every other data processing step performed. BBDuk v38.9 was used to remove adapter sequences and trim sequences to a minimum of 36 bp. Sortmerna v4.3.4 was used to remove rRNA sequences from all samples, including bacterial and eukaryotic rRNA sequences. The logs were compiled using multiqc and checked for quality and to make sure rRNA sequences were completely removed.

Samples prepared in lung tissue were mapped against the pig genome (Sus scrofa: NCBI, GCF000003025.6) using HISAT2 v2.1.0, and the aligned reads were removed from the trimmed reads using Seqtk v1.3-r106 to remove these segments before being mapped against the *P. aeruginosa* genome. All the samples (SCFM and EVPL) were then aligned to the *P. aeruginosa* UCBPP-PA14 genome (NCBI, assembly GCF000014625.1) using HISAT2 v2.1.0 using the BWA v0.7.17-r1188 aligner with the MEM algorithm.

The reads were then mapped and counted using the featureCounts subread package v2.0.3 in R. The reads were mapped against the UCBPP-PA14 strain obtained from Pseudomonas.com.

#### 3.5.6 RNA sequence data analysis

All further analysis took place in Rstudio v2022.07.1 (R Core Team, 2021). Differential gene expression analysis was conducted using DESeq2 v3.15 (Love MI, 2014) on the counted reads. Here differential expression was performed between the different growth environments for each PA14 genotype. A gene was considered statistically significantly differentially expressed if it had a P value ≤ 0.05 (Benjamini-Hochberg procedure to control false discovery rate) and a Log_2_FC of ≥ 1.5. Principal Component Analysis (PCA) was performed using DESeq2 data with the built in ‘plotPCA’ function in the DESeq2 package and 95% confidence ellipses were determined using the ‘stat ellipsis’ function from ggplot v3.3.6 (Wickham, 2016). PCA data were plotted using ggplot. Venn diagrams were created using the VennDiagram package in R. These compared the effect of strain on gene expression in the EVPL compared with growth in SCFM, and compared the effect of treatment on the strains when grown in the EVPL environment.

Kyoto Encyclopaedia of Genes and Genomes (KEGG) (Kanehisa, 2002) enrichment analysis was performed to find the pathways which were enriched between the samples using the KEGG code pau for *P. aeruginosa*. This analysis was done using the EnrichKEGG function from the clusterprofiler package. A KEGG pathway was considered to be significantly enriched if it had an adjusted P value *<*0.05 (Benjamini-Hochberg ).

Gene ontology (GO) analysis performed using the r package ‘TopGo’ and a Fisher’s exact test with P value *<*0.05, here the differential expression analysis dataframe was compared with GO ontology terms from pseudomonas.com version 21.1 (Winsor et al., 2016).

Antibiotic resistance prediction analysis used the Comprehensive Antibiotic Resistance Database (CARD) from pseudomonas.com version 21.1 (Winsor et al., 2016). Predicted antimicrobial resistance genes from the database were compared against the list of loci found to be significantly differentially expressed between our growth conditions. Various databases of antibiotic resistance genes (ARGs) exist and have different merits (Doyle et al., 2020); we chose CARD to provide consistency with our previous transcriptomic work on *P. aeruginosa*.

## 4 Results

### 4.1 Imaging of wild-type and biofilm mutants of *P. aeruginosa* in a high-validity model of CF lung infection

Light microscopy, scanning electron microscopy (SEM) and cryogenic scanning electron microscopy (CryoSEM) were used to image the populations that formed on the porcine tissue after two days of growth. Clear images of bacteria were obtained for tissue colonised by PA14 WT and *pelA^−^*. Haematoxylin-eosin stain cross-sections of PA14 WT (Figure 1A, left column) show a thick layer of biofilm on the epithelial surface of the lung sections, consistent with previous work showing that two days is a sufficient amount of time for a mature biofilm to form (Harrington et al., 2020). Comparable cross-sections of the *pelA*^-^ mutant show either a thin layer of bacteria that has detached from the tissue during processing, or isolated patches of bacteria at the epithelial surface (Figure 1B, left column). The SEM images of the surface of the WT tissue-associated bacterial populations (Figure 1A, centre and right columns) show large numbers of bacteria growing on top of the collagen fibrils of the lung. The SEM images of *pelA^−^* (Figure 1B, centre and right columns) also show the presence of a large number of bacteria, despite the thinness of this layer when viewed in cross-section under a light microscope. This is consistent with our previous work showing that this mutant does grow over the surface of the EVPL tissue and proliferate to high cell density, but consistent with the literature on the effect of pel loss (Colvin et al., 2011), this layer is thin, unstructured and lacking in matrix polysaccharide. We direct the reader to Figs 3,4 & S9 in Harrington et al., (2020) for comparison. The use of CryoSEM retains more of the highly hydrated biofilm matrix and tissue structure, allowing us to determine whether the populations seen in Figure 1 are truly biofilms (Bakker, 2026). The extracellular matrix of bronchioles is primarily composed of collagen and elastin, and the laced structure of collagen fibrils and elastin fibrils in the lung, on which *P. aeruginosa* proliferates and forms biofilms, is more clearly seen in CryoSEM images of uninfected EVPL tissue incubated in SCFM for two days (Figure S1). CryoSEM shows the WT PA14 biofilm in greater detail, with an extensive matrix and a honeycomb structure of matrix-embedded cells punctuated by gaps (Figure 2 A-D). In Figure 2C, there are cells woven and embedded within the porcine tissue itself, with the collagen matrix of the bronchi providing a supporting structure for biofilm formation. Figure 2D shows the biofilm forming around the collagen fibrils, leaving void structures where the collagen fibrils are less densely packed together and make way for the elastin fibrils. In Figure 2D, parts of the biofilm structure surrounding the cells and adhering them together can clearly be seen. We were unable to gain CryoSEM images of *pelA^−^-*colonised tissue, because the bacterial populations detached from the samples during preparation (presumably during freezing). This is consistent with reduced attachment of the mutant populations to one another and the lung tissue, providing further evidence for the lack of mature matrix.

**Figure 1:**
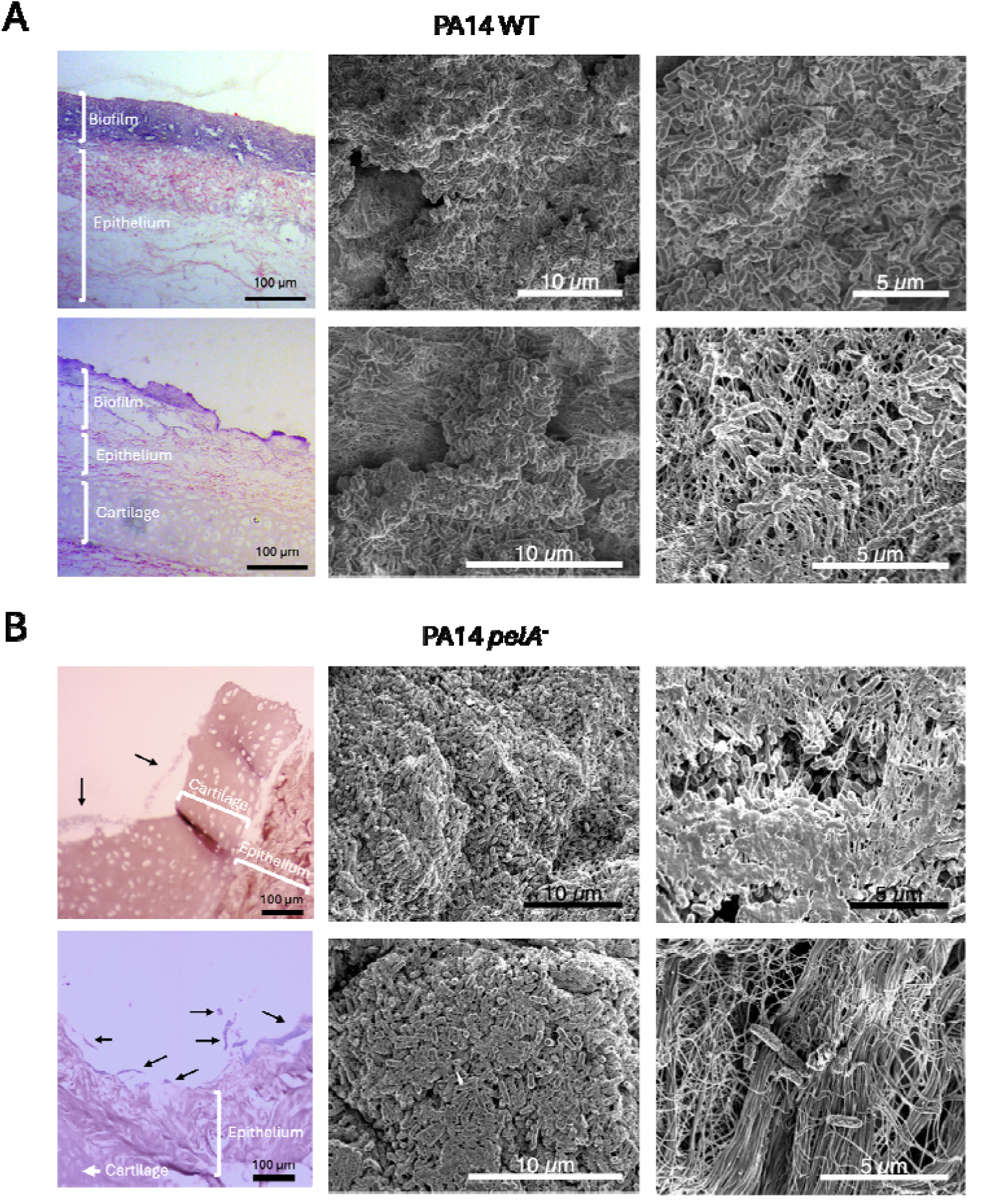
Light and scanning electron microscopy of *P. aeruginosa* PA14 WT (A) and *pelA^−^*(B) grown in the EVPL model for 48h. (A) WT. Light microscopy of two example regions of cross-sectioned tissue (left column) stained with haematoxylin-eosin show thick, structured biofilm (purple) overlying damaged epithelial tissue on top of the basal cartilage of the bronchiole. SEM (centre and right columns) shows large aggregates of cells with some signs of matrix. (B) *pelA*^-^ mutant. Light microscopy of two example regions of cross-sectioned tissue (left column) stained with haematoxylin-eosin show either (top) a thin continuous layer of bacteria (black arrows) or (bottom) small patches of bacteria (black arrows). SEM (centre and right columns) shows the layer in surface view. Light microscopy were images obtained with a Zeiss AxioScope.A1. The two genotypes are shown at slightly different magnifications to allow the respective fields of view to capture key features of both populations. SEM images were obtained using a Zeiss Gemini electron microscope with an InLens detector, at 1kV.

**Figure 2:**
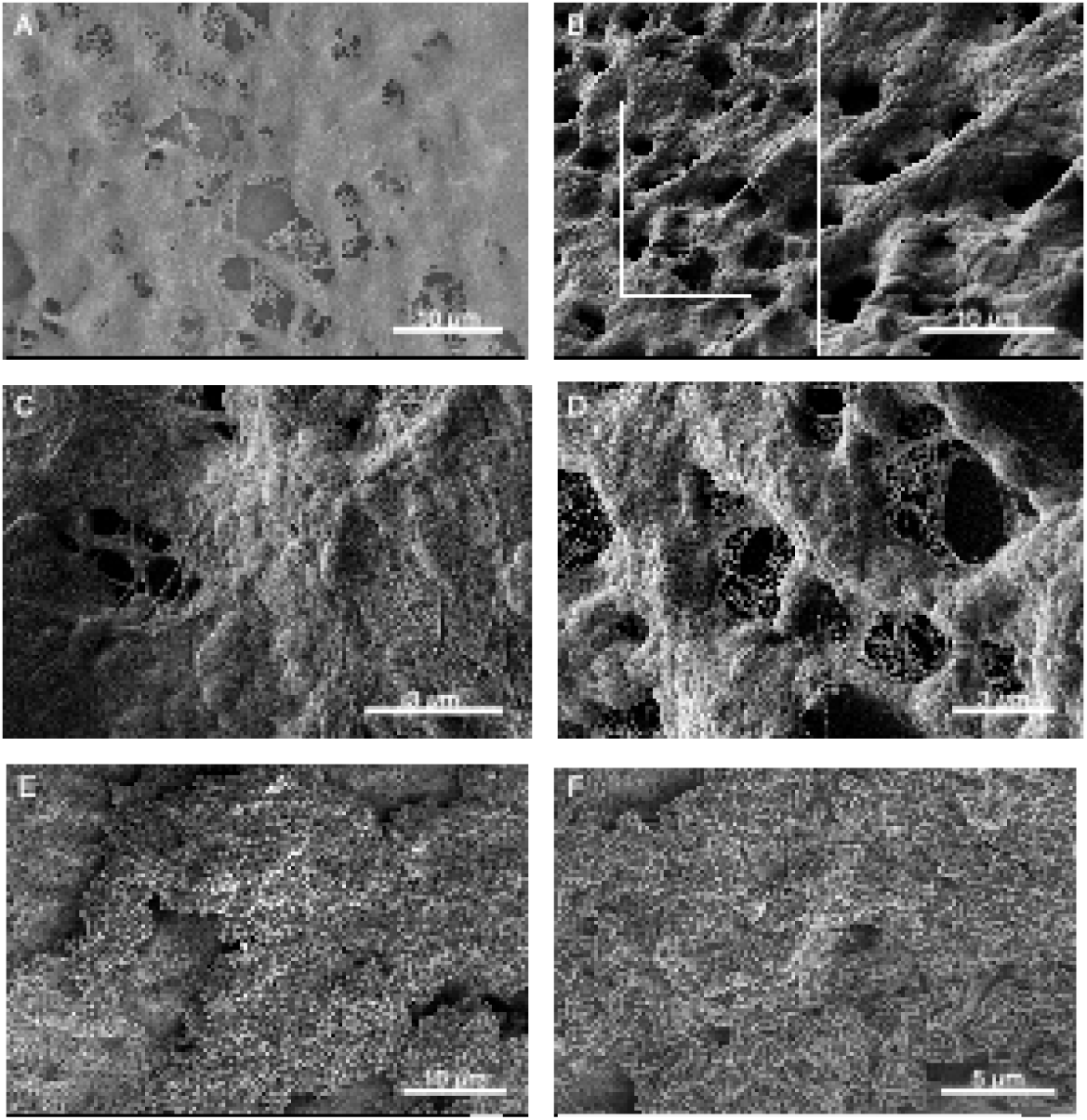
Cryogenic scanning electron microscopy (A-D) and scanning electron microscopy (E-F) images of *P. aeruginosa* strain PA14 grown on EVPL model for 48 hours. Cryogenic scanning electron microscopy images obtained using a Zeiss Crossbeam 550 SEM equipped with a quorum 3010 Cryo-system performed at 2 kV. Scanning electron microscopy images obtained using a Zeiss Gemini electron microscope with an InLens detector, at 1kV.

**Figure 3:**
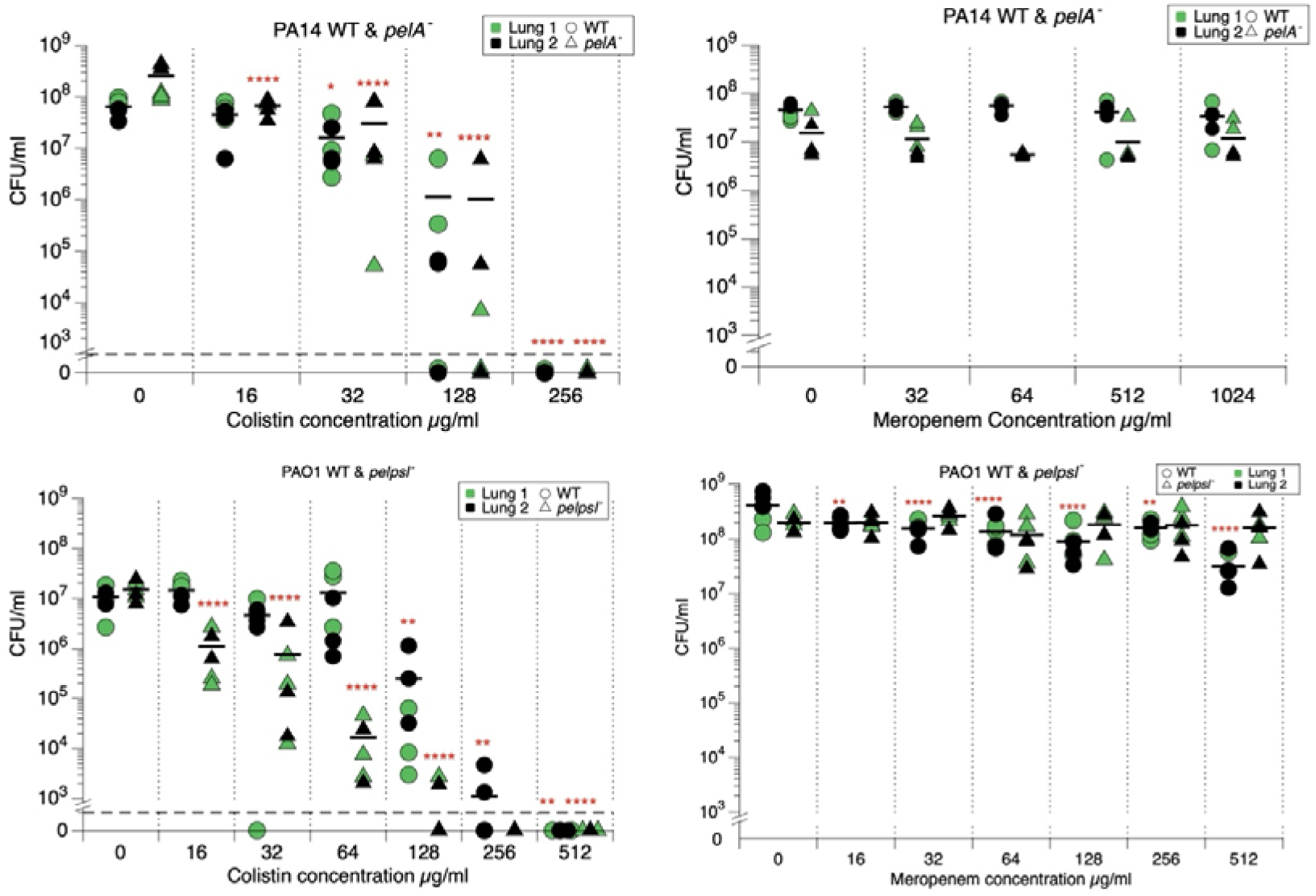
Antimicrobial susceptibility testing in EVPL model of WT and biofilm mutants of *P. aeruginosa* strains PA14 and PAO1. Each symbol represents a single piece of lung tissue, with tissue from two different pigs shown in black/green. Black bars show the mean CFU/ml of lung 1 and lung 2 per condition. ANOVA or multiple Mann Whitney with post-hoc Dunnett’s or Dunn’s test were used for comparison of treated conditions with the untreated control, t-tests found no differences between WT and mutants at each antibiotic concentration. The red asterisk (*) denotes a statistically significant difference from the untreated control. *= ≤ 0.05, **= *<*0.01, *** = *<*0.001 and **** = ≤0.0001. P values were corrected with False discovery rate (FDR). PA14 colistin: genotype*treatment F_4, 50_ = 6.27 P=0.0004, treatment F_4, 50_ = 16.57 P<0.0001, genotype F_1,50_ = 9.62 P=0.0032. PAO1 colistin: interaction F_6, 69_ = 5.78 P < 0.0001, treatment F_6, 69_ = 11.90 P<0.0001, genotype F_1, 69_ = 11.90 P=0.001. PA14 meropenem: genotype*treatment F_4,50_ = 0.857 P=0.496, treatment F_4, 50_ =0.538 P=0.708, genotype F_1, 50_ = 9.95 P=0.0027. PAO1 meropenem: genotype*treatment F_6, 70_ = 4.49 P=0.0007, treatment F_6, 70_ = 6.28 P<0.0001, genotype F_1, 70_ = 0.697 P=0.4068. Raw data is provided in the Data Supplement.

**Figure 4:**
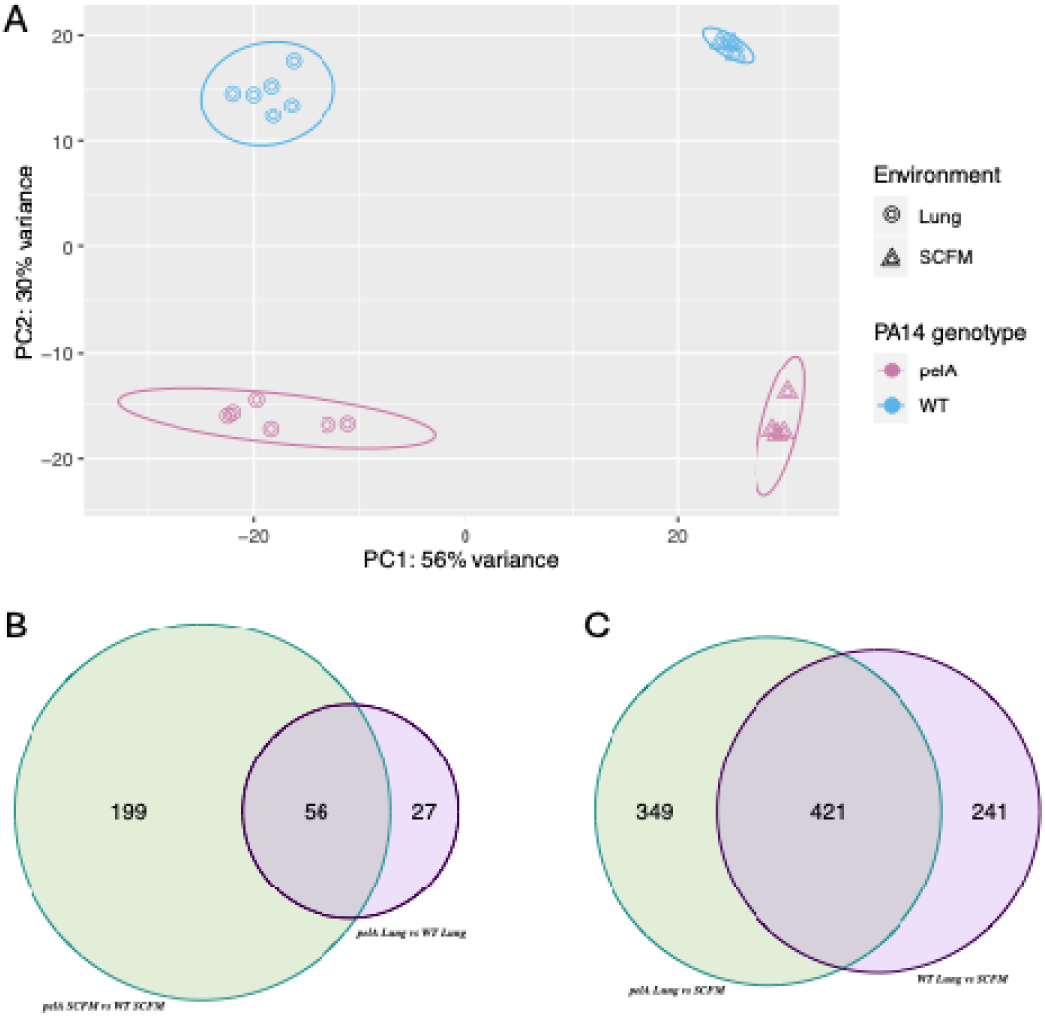
Principal component analysis (PCA) and Venn diagram of differential gene expression by *P. aeruginosa* PA14 WT and *pelA^−^*strains grown on the EVPL model or grown in SCFM only. (A) PCA plt with 95 % confidence ellipsis produced by the stat ellipsis function in ggplot2. (B,C)Venn diagrams of differentially expressed genes (DEGs). (B) The effect of strain and growth environment, *pelA^−^* and WT grown in the **EVPL** vs SCFM. (C) Shared and unique genes between WT **EVPL** vs SCFM and *pelA^−^* EVPL vs SCFM. Gene expression threshold was set at P value 0.05 and |LFC| 1.5. The shared DEGs are those which are either upregulated or downregulated in both treated *pelA^−^* and WT genotypes compared with their untreated counterparts.

Having confirmed the biofilm defect of the PA14 *pelA*^-^ mutant, and the reproducibility of the contrast between the WT and mutant reported in previous research, we thus have a strong foundation for novel work that compares matrix mutants to the WT in terms of antibiotic tolerance and transcriptome.

### 4.2 Biofilm mutants of *P. aeruginosa* do not show reduced tolerance to colistin or meropenem in a high-validity model of CF lung infection

*P. aeruginosa* populations formed after two days of growth in the EVPL model were then treated with colistin or meropenem. The lung pieces were transferred to fresh SCFM containing a concentration series of either antibiotic, or to fresh SCFM containing no antibiotics, and incubated for a further 24 h before recovery of bacteria for plating. Plate counts of viable bacterial load associated with the tissue confirmed that, consistent with previous work (Harrington et al., 2020), both PA14 WT and PA14 *pelA^−^* grew to similarly high densities in the model (10^7^-10^9^ CFU ml^-1^, Figure 3). Contrary to our expectations, loss of pel did not lead to a reduction in tolerance to colistin or meropenem when grown in the EVPL model. As shown in Figure 3, the PA14 *pelA^−^* mutant and the WT were able to survive at the same concentrations. To determine whether this effect was specific to PA14, which does not produce psl, we repeated this experiment using the psl-expressing strain PAO1 and an isogenic double *pel^−^psl^−^* mutant. The same result was observed: the mutant lacking both structural biofilm exopolysaccharides was just as tolerant of colistin and meropenem as the WT (Figure 3). Several explanations for this are possible. First, the mutant may compensate for lack of a protective matrix by quantitatively or qualitatively different expression of resistance determinants than the wild type (e.g. increased production of efflux pumps, or expression of a more effective efflux pump). Second, the model growth conditions may enforce bacterial aggregation regardless of matrix production, leading to similar tolerance levels (Goltermann and Tolker-Nielsen, 2017).

### 4.3 Transcriptomic analysis of *P. aeruginosa* PA14 WT and the *pelA* mutant grown in the EVPL model and SCFM *in vitro*

We previously established that growth of PA14 WT in the EVPL model cued increased expression of certain antimicrobial resistance (AMR) determinants compared with growth in SCFM *in vitro*, in the absence of antibiotic treatment to act as a specific cue for these genes (upregulation of *arnA*, *crpP*, downregulation of the efflux pump repressor *mexL*: (Harrington et al., 2022)). We therefore next asked whether the *pelA^−^*mutant increased expression of the same AMR genes as the WT, to the same extent, when grown in the EVPL model. We hypothesised that the mutant might show enhanced upregulation of these loci, and/or promote expression at loci not upregulated in the WT, to compensate for a lack of a matrix. In other words, did the environment in the EVPL cause genotype-independent induction of a phenotype so antibiotic tolerant that any additional protection conferred by the matrix became redundant, or did the mutant use a different physiological response to confer protection in the absence of a matrix?

We performed RNA sequencing (RNAseq) to ascertain the transcriptome of PA14 WT and *pelA^−^* (a) in SCFM *in vitro*, in the absence of antibiotics; (b) in the EVPL model, in the absence of antibiotics; and (c) in the EVPL model, in the presence of sub-bactericidal colistin. Due to financial constraints, we focused on just colistin as we had previously assessed penetration of this molecule into the biofilm matrix in the EVPL model. By comparing expression between genotypes and conditions, we were able to assess differences in responses to SCFM, to the EVPL, and to colistin in the EVPL model.

PA14 WT and *pelA^−^* were grown for 48 h either in the EVPL model, or in SCFM alone in static culture plates. In the latter condition, PA14 WT aggregated and formed a pellicle. Figure S2 shows the difference in biofilm formation by the two genotypes in this *in vitro* growth platform. Replica sets of 48 h EVPL biofilms were treated with either 64 µg/ml colistin, or an equivalent volume of SCFM. First, we compared the gene expression of each genotype when grown in the EVPL versus in SCFM *in vitro*, in the absence of colistin as a control. Second, we compared expression of each genotype to growth in sub-bactericidal colistin, (as determined by antimicrobial susceptibility testing in the EVPL, Figure 3), versus mock treatment in the EVPL. Gene expression was considered significantly different when the log_2_ fold-change (LFC) was both statistically significantly different from 0 (P value corrected for multiple testing with Benjamini and Hochberg method <0.05), and had a value ≥1.5 or ≤-1.5.

Principal component analysis (PCA) was first used to assess the variance in the gene expression data (all genes, n=5289) between genotypes and growth environments in the absence of antibiotic treatment. As shown in Figure 4, there were significant differences in the transcriptome between genotypes and growth environments. Growth environment was found to be the biggest driver of variance, consistent with our previous work.

### 4.4 Overview of differentially-expressed genes in EVPL vs. SCFM for WT and *pelA*-mutant in the absence of colistin

Kyoto Encyclopaedia of Genes and Genomes (KEGG) analysis was used to show the pathways that were enriched between growth in EVPL vs SCFM (Figure S3). For PA14 WT, there were eleven significantly enriched pathways in EVPL vs SCFM. Nine of these pathways were associated with metabolism such as pyruvate and nitrogen metabolism, and multiple pathways associated with fatty acid metabolism. The phenazine and quorum sensing pathways were also enriched, with all or most (respectively) genes associated with these pathways downregulated. This is consistent with our previous study comparing the transcriptome of PA14 WT in the EVPL vs. SCFM (Harrington et al., 2022). In comparison, *pelA*^−^ only had four enriched pathways. Out of these four pathways, two are shared with WT: phenazines and quorum sensing. The phenazine pathway and the majority of quorum sensing loci were also downregulated in *pelA*^−^ when grown in EVPL vs SCFM. This further highlights the more chronic-like infection phenotype when *P. aeruginosa is* grown in EVPL vs SCFM. The two enriched pathways unique to the mutant were flagellar assembly and bacterial chemotaxis, both of which were downregulated in EVPL vs. SCFM. KEGG analysis comparing *pelA^-^* vs. WT transcriptomes in both the EVPL and SCFM showed downregulation of two-component systems *(pctA* and *pctB*, which are chemoreceptors involved in chemotaxis towards amino acids) flagellar assembly and bacterial chemotaxis in the mutant, compared with the WT (Figure S3).

More detailed gene ontology (GO) enrichment analysis found a number of shared enriched processes when comparing growth of PA14 WT and *pelA****^−^*** in EVPL vs SCFM (Figure S4, S5). These include negative regulation of transport and protein secretion and type IV pilus system. The processes which were enriched only in PA14 WT included metabolic and catabolic processes such as lipid metabolism, glutamine biosynthetic process and fatty acid metabolism and biosynthesis. The processes enriched only in *pelA****^−^*** were flagellar assembly, transport of cations and siderophores, chemotaxis and antibiotic biosynthetic process (this last GO term includes loci for phenazine biosynthetic and enterobactin-mediated iron transport). The results of the GO analysis were thus broadly consistent with the KEGG analysis, and did not immediately present any unique response to the EVPL by the *pelA-* mutant that could protect it from antibiotics in the absence of biofilm matrix.

### 4.5 The *pelA* mutant does not express higher levels of canonical resistance determinants in response to the EVPL environment

It is possible that the *pelA^−^* mutant responded to environmental cues in SCFM and/or the EVPL tissue by switching on a different set of specific antibiotic resistance determinants that ‘prime’ it to survive later antibiotic exposure, compared with the WT, leading to the phenotype observed in Figure 3. KEGG classifies many pathways into very broad groups. For instance, the KEGG pathway for cationic antimicrobial peptide resistance includes modifications to the inner membrane, outer membrane and peptidoglycan, as well as mechanisms for both intracellular and extracellular degradation of antimicrobial peptides. So if only one of these mechanisms is upregulated in response to colistin administration, it would be unlikely to result in the whole KEGG group being significantly enriched. We therefore followed a more targeted approach by using the CARD (Alcock et al., 2023) database to focus on differential expression on specific loci and operons that are associated with AMR.

A number of efflux pumps were discovered to be significantly up or down regulated in at least one of the conditions. However, the CARD analysis did not show any significantly differentially expressed genes (DEGs) between *pelA^−^*vs WT grown in the EVPL, or *pelA^−^* vs WT in SCFM. i.e. this analysis suggested that both genotypes showed similar up- or down-regulation when growth in the EVPL was compared with growth in SCFM *in vitro*.

However, the CARD analysis did show the importance of efflux pumps in the increase of antibiotic tolerance, in particular the resistance nodule division (RND) family, in both genotypes (Figure 5). These are multidrug efflux pumps responsible for increased innate resistance in *P. aeruginosa*. The RND efflux pumps have been noted as one of the most important systems for multidrug resistance (Scoffone et al., 2021). Interestingly the MexAB-OprM pump repressor gene *nalD* is upregulated in the EVPL vs SCFM in both genotypes, and the efflux pump genes are also downregulated in both. This efflux pump is known to confer resistance against some carbapenems including meropenem (Atrissi et al., 2021). Similar changes in *nalD* and *mexAB-oprM* expression were observed for PA14 WT in EVPL vs. SCFM in our previous study, although they were not found to be significant in that analysis (Harrington et al., 2022). Interestingly, other efflux pumps were upregulated in EVPL vs SCFM in both genotypes, with only the *oprM* porin downregulated (see Figure 5).

**Figure 5:**
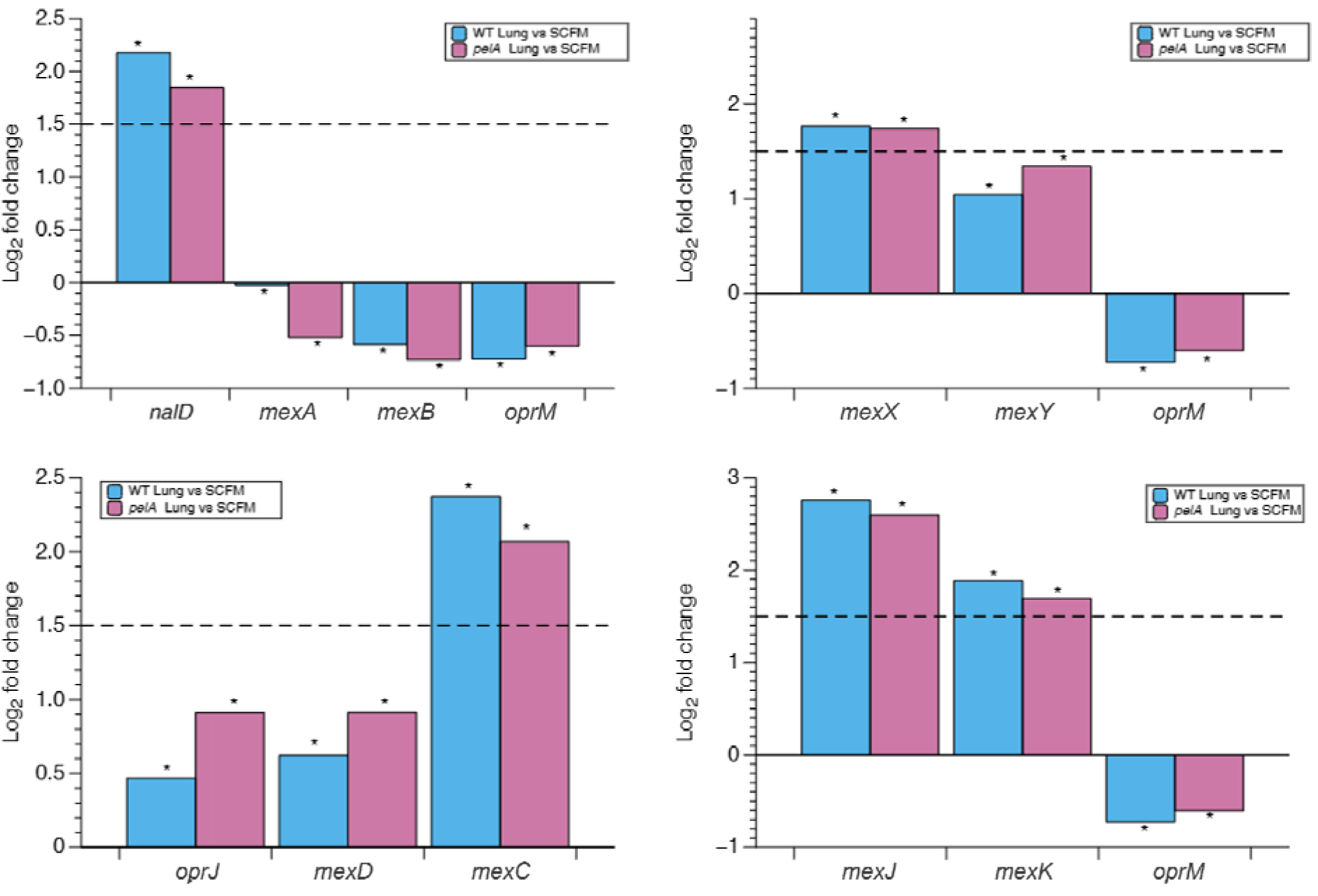
The log_2_ fold change (LFC) in expression of genes involved in efflux pump systems when *P. aeruginosa* PA14 WT and *pelA^-^* were grown in the *ex vivo* pig lung (EVPL) vs synthetic cystic fibrosis sputum media (SCFM) *in vitro*. These genes of interest were found in the Comprehensive Antibiotic Resistance Database (CARD). Gene expression analysis was performed between PA14 WT and *pelA^−^* grown for 48 hours in either the EVPL or SCFM environments. The asterisk * denotes an LFC significantly different from zero result where the p_adj_ ≤0.05 The dotted lines denote an LFC of 1.5. Gene expression was considered significantly different when |LFC | was ≥ 1.5 and p_adj_ was ≤0.05. P value corrected for multiple testing with Benjamini and Hochberg method

We also found the TetR family, associated with cell stress response and AMR, transcriptional regulator *tetR* to be significantly upregulated in the EVPL vs SCFM in both WT and *pelA^−^* as, seen in Figure S6. As well as other genes associated with carbapenem resistance, including the porins *oprD* and *opdP*, and the beta-lactamase gene *ampC* were upregulated (Rostami et al., 2018). The log2-fold changes in expression of *oprD*, *opdP* and *ampC* are shown in Figure 6.

**Figure 6:**
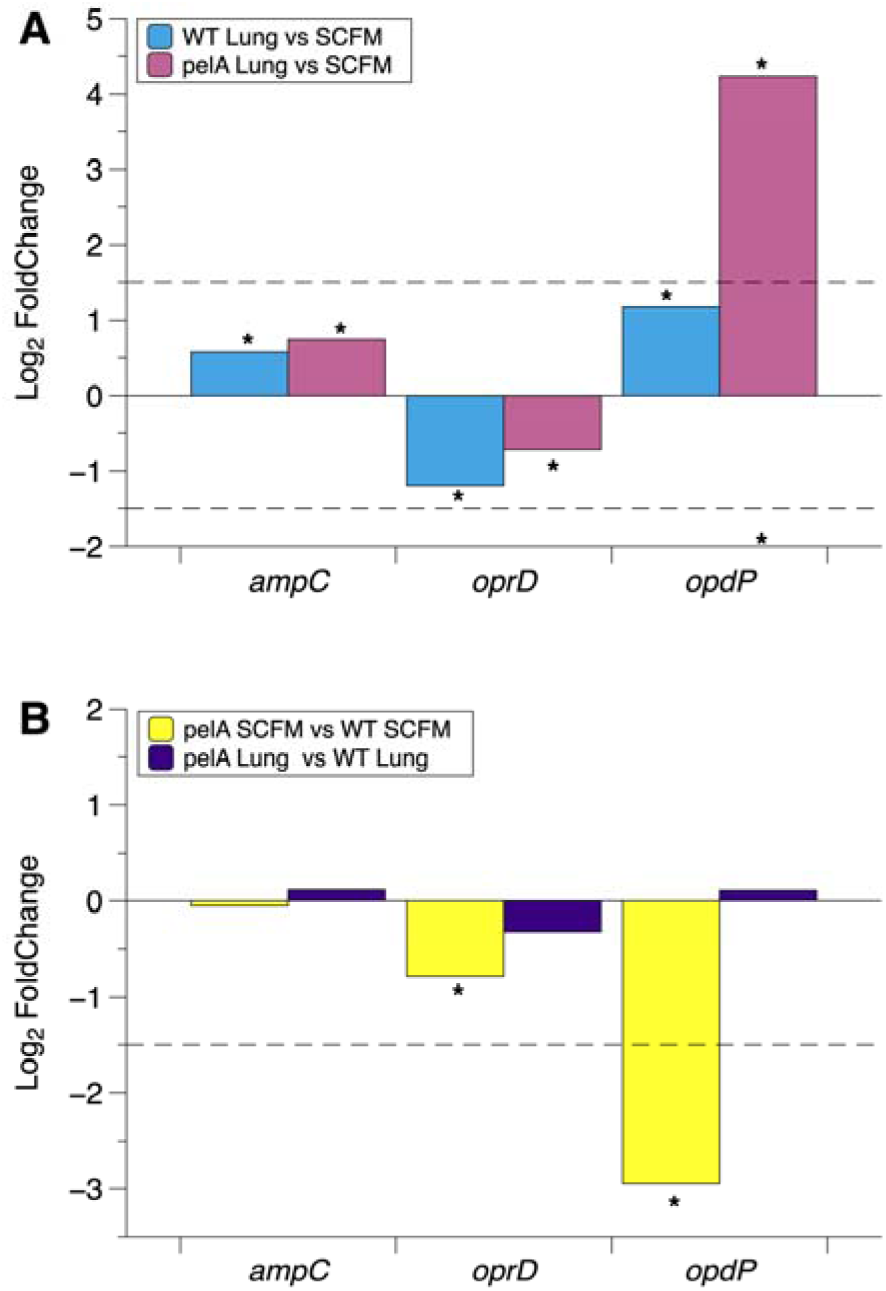
Log_2_ FoldChange (LFC) in expression of carbapenem resistance genes found in *P. aeruginosa* PA14 WT and *pelA^−^* when comparing growth in the *ex vivo* pig lung model (EVPL) and synthetic cystic fibrosis sputum media (SCFM) *in vitro*. These genes of interest were found in CARD. (A) Comparisons between growth in the EVPL model vs SCFM for both genotypes. (B) Comparisons between genotypes when grown in SCFM and the EVPL model. Black dotted lines denote an LFC of 1.5 or –1.5. The asterisk * denotes an LFC significantly different from zero result where the p_adj_ ≤0.05 . Gene expression was considered significantly different when |LFC| was ≥ 1.5 and p_adj_ was ≤0.05. P value corrected for multiple testing with Benjamini and Hochberg method.

Though significantly different, the change in expression of the beta-lactamase *ampC* gene in EVPL vs SCFM did not reach the Log2 fold change cutoff for either WT or *pelA^−^* (Figure 6A). There was no significant difference found in expression levels of *ampC* when comparing WT vs *pelA^−^* in either the EVPL or SCFM, suggesting that this mechanism is not induced by growth in either environment (Figure 6B). The gene *oprD* was downregulated, though did not reach our log2 fold cutoff, in the EVPL vs SCFM in both genotypes with slightly less expression in *pelA^−^* vs WT for both environments. Though not significantly downregulated, a reduction in this porin is associated with higher resistance to carbapenems as it allows entry of meropenem into the cell. The lower expression levels of *oprD* in the EVPL compared with SCFM may explain higher levels of tolerance to meropenem in the EVPL. Interestingly, *opdP* was significantly upregulated in the EVPL vs SCFM for *pelA^−^* but showed no significant change for the WT. However, expression of *opdP* was significantly lower for *pelA^−^* vs WT when grown in SCFM, and the greater increase in expression by the mutant in EVPL resulted in expression levels that were similar to the WT when grown in the EVPL (Figure 6B).

*P. aeruginosa* exhibits resistance to cationic polypeptide antibiotics, such as colistin, through a variety of mechanisms. One of the major mechanisms is the *arn* operon (Janet-Maitre et al., 2024), which acts to change the charge of the lipid A molecule on the LPS to which colistin binds, reducing its efficacy. As observed in Figure 7 there are differences in expression of the operon in SCFM vs. EVPL for each genotype, and between genotypes in each growth condition.Three genes were found to be significantly upregulated by *pelA^−^*in the EVPL vs. SCFM: *arnE*, *arnF* and *arnT*. These genes were also upregulated in the same comparison for the WT, but did not reach the significant LFC threshold of ≥ 1.5. Whilst there was upregulation of these genes in EVPL vs SCFM there was still less expression in the *pelA^−^* mutants than in WT.

**Figure 7.**
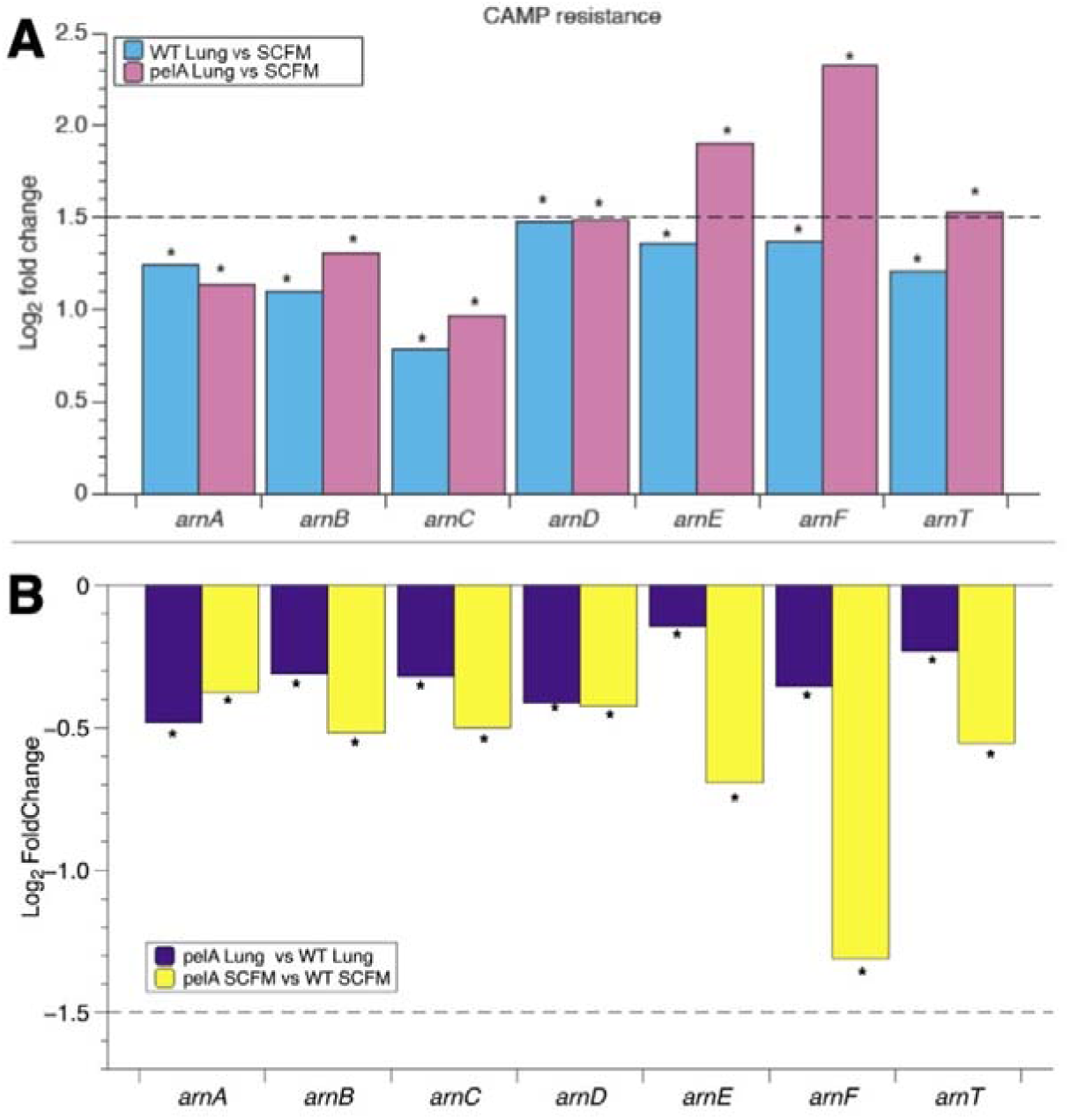
Log_2_ FoldChange (LFC) in expression of CAMP resistance genes found in *P. aeruginosa* PA14 WT and *pelA^−^* when comparing growth in the *ex vivo* pig lung model (EVPL) and synthetic cystic fibrosis sputum media (SCFM) *in vitro*. These genes were found in CARD. (A) Comparisons between growth in the EVPL model vs SCFM for both genotypes. (B) Comparisons between genotypes when grown in SCFM and the EVPL model. Black dotted lines denotes a LFC of 1.5 or –1.5. The asterisk * denotes an LFC significantly different from zero result where the p_adj_ ≤0.05. Gene expression was considered significantly different when |LFC | was ≥ 1.5 and p_adj_ was ≤0.05. P value corrected for multiple testing with Benjamini and Hochberg method.

Overall, there was very little difference in expression of canonical AMR genes between the two genotypes when grown in the EVPL model in the absence of any antibiotics.

### 4.6 Alginate did not compensate for lack of pel polysaccharide

We next investigated whether *pelA^−^* upregulated expression of alginate biosynthetic genes more than the WT, in order to compensate for the lack of the pel polysaccharide. Although alginate is not a major structural polysaccharide in *P. aeruginosa*, it helps adhesion to surfaces, and its overproduction can enhance antibiotic tolerance (Wozniak et al., 2003).

There was no significant difference in expression for 12 out of the 13 genes involved in the production of alginate in any of the comparisons, as shown in TableS1. The gene *algD* was significantly upregulated in the WT grown on the EVPL vs SCFM, but expression was similar between EVPL and SCFM for *pelA^−^*. Interestingly, there was actually lower expression of the alginate biosynthesis genes for *pelA^−^* compared with WT for both EVPL and SCFM, although these differences were not statistically significant. These results corroborate results from other studies which suggest that alginate is not sufficient to replace the pel polysaccharide (Wozniak et al., 2003). This is consistent with the crystal violet staining results shown in Figure S2.

### 4.7 Overview of differentially-expressed genes in response to colistin in the EVPL for WT and *pelA*- mutant

We next explored transcriptomic responses of the WT and *pelA^−^* mutant to sub-bactericidal concentrations of colistin in the EVPL model, to see if these differed. PCA (Figure 8) showed that gene expression varied more between genotypes than treatment conditions. Just 100 and 105 genes were significantly differentially expressed in response to colistin by either the WT or *pelA^−^*mutant exclusively, respectively. There were 50 genes significantly differentially expressed by both genotypes. The full list of DEGs is provided in the Data Supplement. Although expression of alginate-related genes was not found to differ between the WT and mutant, we checked whether colistin treatment had an effect, as alginate is thought to have protective properties for *P. aeruginosa* biofilms. However, there were no significant differences between gene expression in treated vs untreated samples for both the WT and *pelA^−^* (Table S2). The WT upregulated the expression of all loci in the pel operon in response to colistin, but none of these changes were statistically significant (Figure S7).

**Figure 8.**
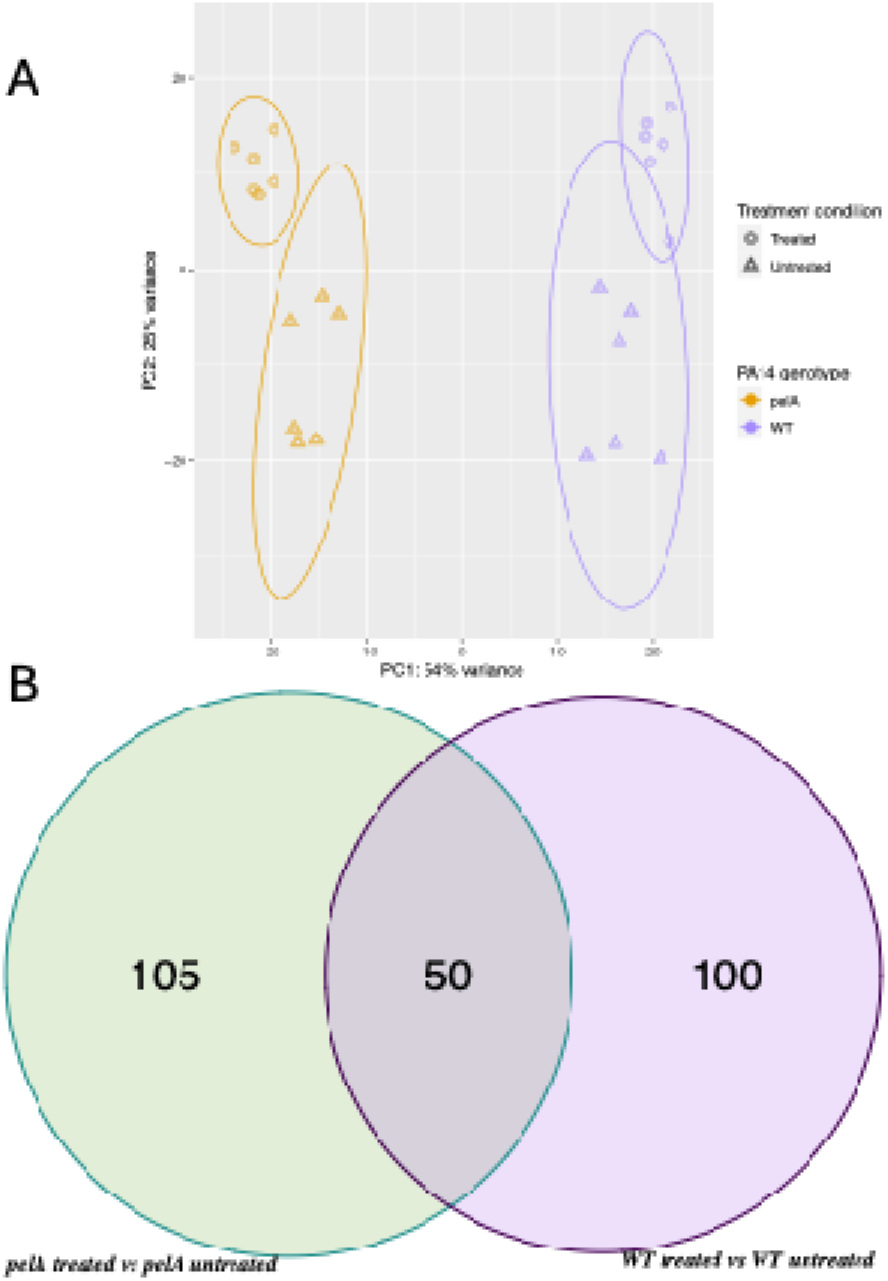
Principal component analysis (PCA) and Venn diagram of differential gene expression by *P. aeruginosa* PA14 WT and *pelA^−^*strains grown on the EVPL model, in the presence vs. absence of colistin. A) PCA plot showing variance of differentially expressed genes (DEGs) from PA14 WT and *pelA^−^* grown on the EVPL model and either left untreated or treated with colistin, 95 % confidence ellipsis have been produced by the stat ellipsis function in ggplot2 for the PCA plots. (B) Venn diagrams are of differentially expressed genes (DEGs). Venn diagram of shared and unique significant differentially expressed genes between two contrasts, *pelA^−^* treated vs untreated and WT treated vs untreated. Gene expression threshold was set at p_adj_ value of ≤0.05 and |LFC| of ≥ 1.5.

KEGG analysis revealed four enriched pathways in both WT and the *pelA*^-^ mutant when comparing colistin treated vs. untreated conditions (Figure 9). Three of these - nitrogen metabolism (all downregulated), two component systems (including both up- and down-regulated loci) and CAMP resistance (all upregulated) – were common to both genotypes. In the WT, amino sugar and nucleotide metabolism was also enriched (all upregulated), while in the *pelA*^-^ mutant the valine, leucine and isoleucine degradation pathway was enriched (all downregulated).

**Figure 9:**
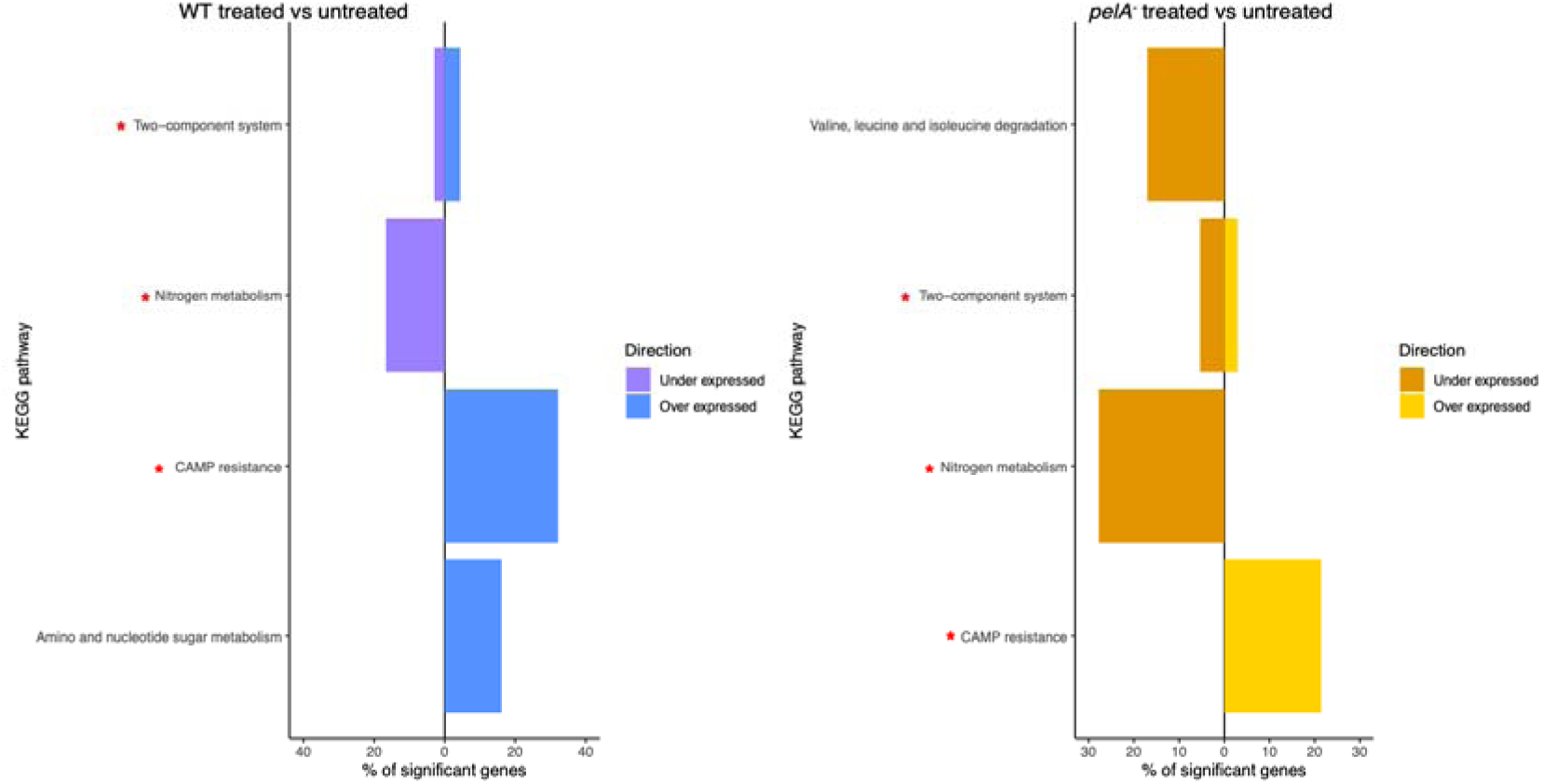
KEGG analysis of enriched pathways in *P. aeruginosa* PA14 WT and *pelA^-^* grown in the EVPL in the presence vs absence of sub-bactericidal colistin. The bar charts show the percentage of genes that are significantly up- or downregulated in each significantly-enriched pathway. Genes that are upregulated are shown right of the zero line, and genes that are downregulated are shown to the left. Genes were classed as significantly differentially expressed if they had a |Log2 FoldChange| ≥ 1.5 and a p_adj_ < 0.05. The red asterisk show that the pathways are enriched in both genotypes. P-values were corrected for multiple testing with the Benjamini and Hochberg method.

GO analysis (Figure 10) found a number of enriched pathways between treated and untreated samples, some of which are shared by both the WT and the *pelA*^-^ mutant, including nitrate metabolic process (consistent with the KEGG analysis) and xenobiotic transport. Genes involved in xenobiotic transport were upregulated in treated vs. untreated samples for both genotypes, and this likely represents colistin efflux; this GO group includes efflux pumps of the RND family. Uniquely enriched pathways for the WT included cell wall organisation, chaperone-mediated protein folding, pilus organisation and amino & nucleotide sugar metabolism – interestingly, this last group includes loci from the *arn* operon (Table S3). The unique pathways in the *pelA*^-^ mutant included cellular respiration, lipid catabolic process, siderophore transport, iron ion transport, arginine catabolic and metabolic process and DNA-templated transcription initiation.

**Figure 10:**
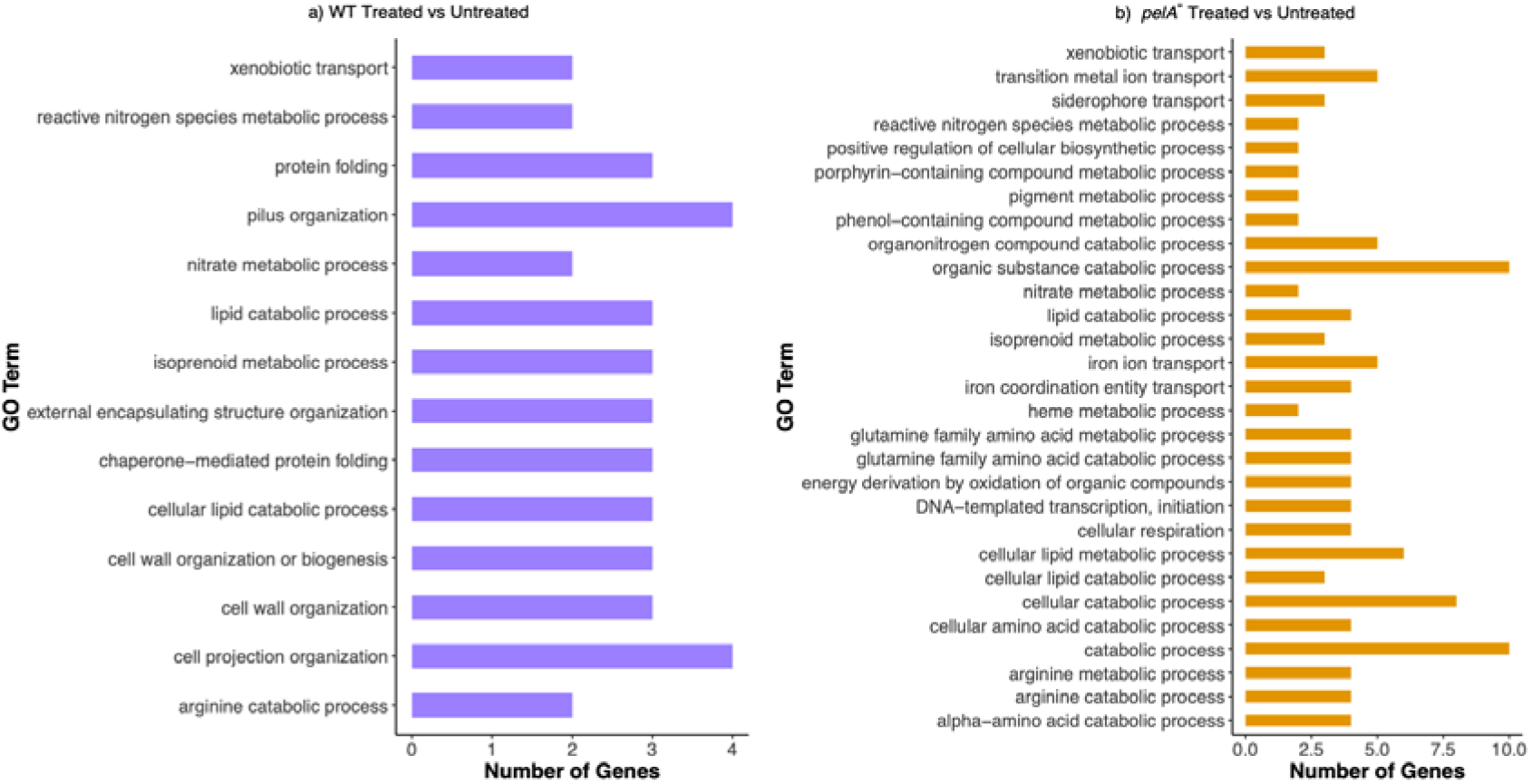
Gene ontology (GO) enrichment analysis of *P. aeruginosa* PA14 (a) WT or (b) *pelA^-^*mutant, grown in the EVPL in the presence vs. absence of sub-bactericidal colistin. The bar graphs show the significantly enriched GO terms (Fisher’s exact test, p < 0.05) with number of DEGs found for each term. Genes were classed as significantly differentially expressed if they had |LFC| ≥ 1.5 and p_adj_ < 0.05. P-values were corrected for multiple testing with the Benjamini and Hochberg method.

### 4.8 Differential expression of MexEF-OprN and cationic antimicrobial peptide resistance expression in WT and *pelA**^−^*** mutant in response to colistin in EVPL

We cross-referenced our lists of genes that were significantly differentially expressed between treated vs untreated comparisons for each genotype with the CARD database to identify predicted ARGs. Efflux pumps of the RND family were predominantly found to be increased in treated vs untreated EVPL samples for both genotypes, but there was a difference in which efflux pumps were differentially expressed in response to colistin (Figure 11). The genes *mexAB-OprM*, *mexXY-OprM* and *mexCD-OprJ* were upregulated in treated vs untreated samples for both WT and *pelA^−^*, but *mexEF-OprN* was upregulated only in the *pelA^−^* mutant.

**Figure 11.**
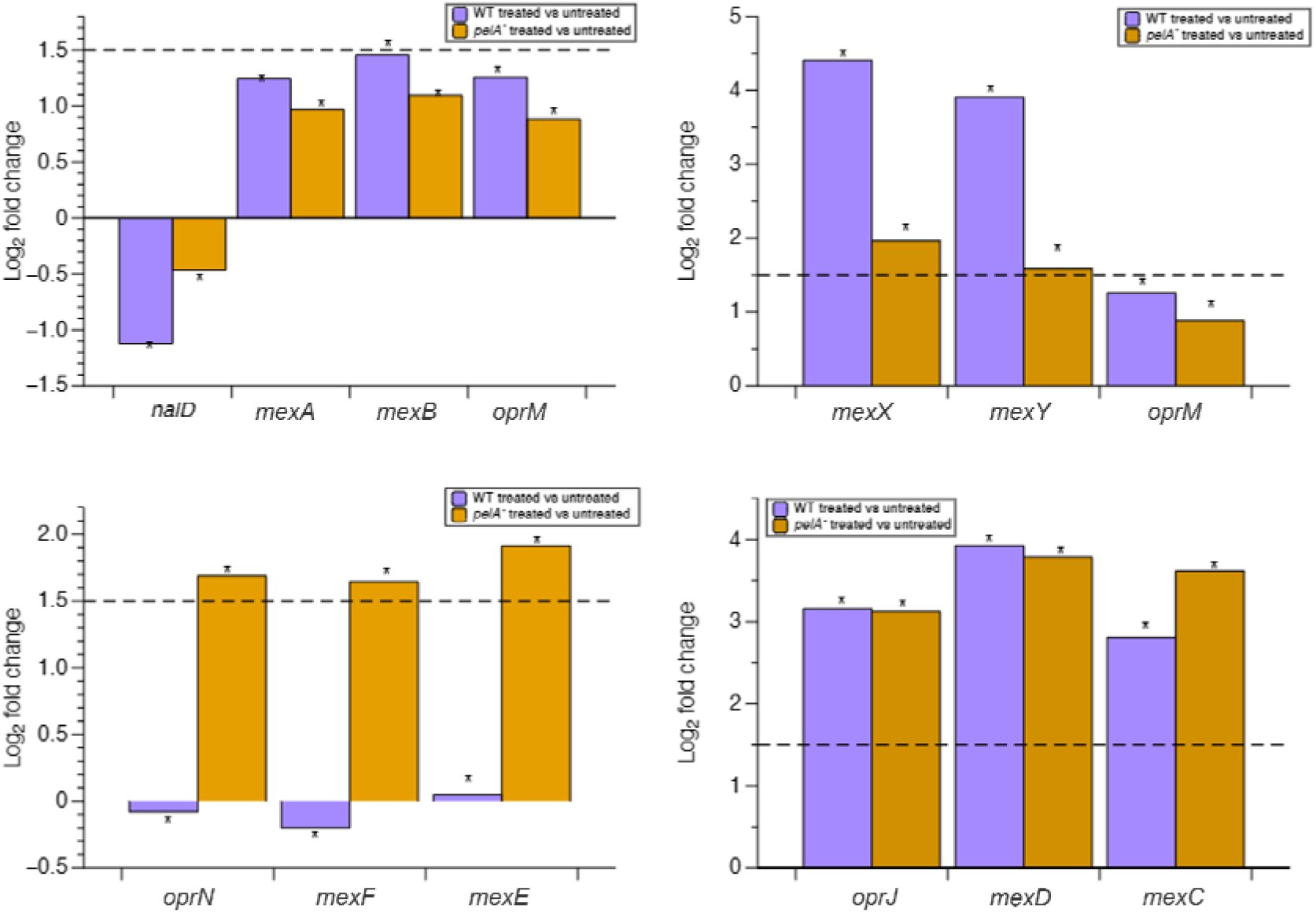
Log_2_ fold change (LFC) in expression of efflux-related genes of interest when *P. aeruginosa* PA14 WT or *pelA^-^* was treated with sub-inhibitory colistin. CARD analysis of genes associated with antibiotic resistance from pseudomonas.com. Gene expression analysis was performed between PA14 WT and *pelA^−^* grown in the EVPL, treated and untreated conditions. The dotted line is an LFC of 1.5. T The asterisk * denotes an LFC significantly different from zero where the p_adj_ ≤0.05 . Gene expression was considered significantly changed when |LFC | was ≥ 1.5 and p_adj_ was ≤0.05. P value corrected for multiple testing with Benjamini and Hochberg method.

The expression of genes involved in the MexAB-OprM pump were found to be downregulated in the EVPL vs SCFM in the absence of colistin, while expression of *nalD*, which suppresses production of this pump, was increased. This efflux pump has been associated with increased resistance to quinolones, β-lactams, tetracyclines and more (Pesingi et al., 2019). It has also previously been shown to be necessary for colistin tolerance in biofilm-associated *P. aeruginosa* PAO1 cells (Pamp et al., 2008a). Consistent with this, in the colistin vs. untreated comparison, we observed a decrease in *nalD* and increases in *mexAB* and *oprM*. However, these changes in expression did not meet the 1.5 LFC benchmark for significance and the pattern of transcriptional change was the same for both the WT and the *pelA^−^* mutant.

Similarly, *mexXY-oprM* and *mexCD-oprJ* were upregulated in the treated vs untreated samples for both genotypes. MexXY-OprM has been associated with resistance to aminoglycoside antibiotics and MexCD-OprJ has been linked with fluoroquinolone resistance (Hocquet et al., 2003, Hirai et al., 1987). Significant upregulation of *mexX* and *mexY* was seen in the WT, and *mexX* was among the top five upregulated genes with the greatest LFC when treated with colistin vs. untreated. Whilst there was also significant upregulation of *mexX* and *mexY* by the *pelA^−^* mutant when treated with colistin, the LFC was smaller than the WT (LFCs: X & Y respectively). Differential expression of genes involved in the MexCD-OprJ pump was comparable in both the WT and *pelA^−^* mutant.

The MexEF-OprN efflux pump, however, was only upregulated by the *pelA^−^*genotype. The MexEF-OprN pump is predominantly associated with increased resistance to fluoroquinolones such as ciprofloxacin (Llanes et al., 2011). It is not typically expressed in WT strains (Llanes et al., 2011), which correlates with the results shown in Figure 9. Interestingly, the MexEF-OprN transcriptional regulator, *mexT,* was not significantly differentially expressed in either PA14 WT (LFC= -0.23) or *pelA^−^* (LFC=-0.05) treatment contrasts.

However, all genes in the *arn* operon, associated with CAMP resistance, were significantly upregulated in the WT, and all but *arnF* and *arnT* were significantly upregulated in the *pelA^−^*mutant (Figure 12). There was a greater LFC observed for the WT than the mutant for all genes.

**Figure 12.**
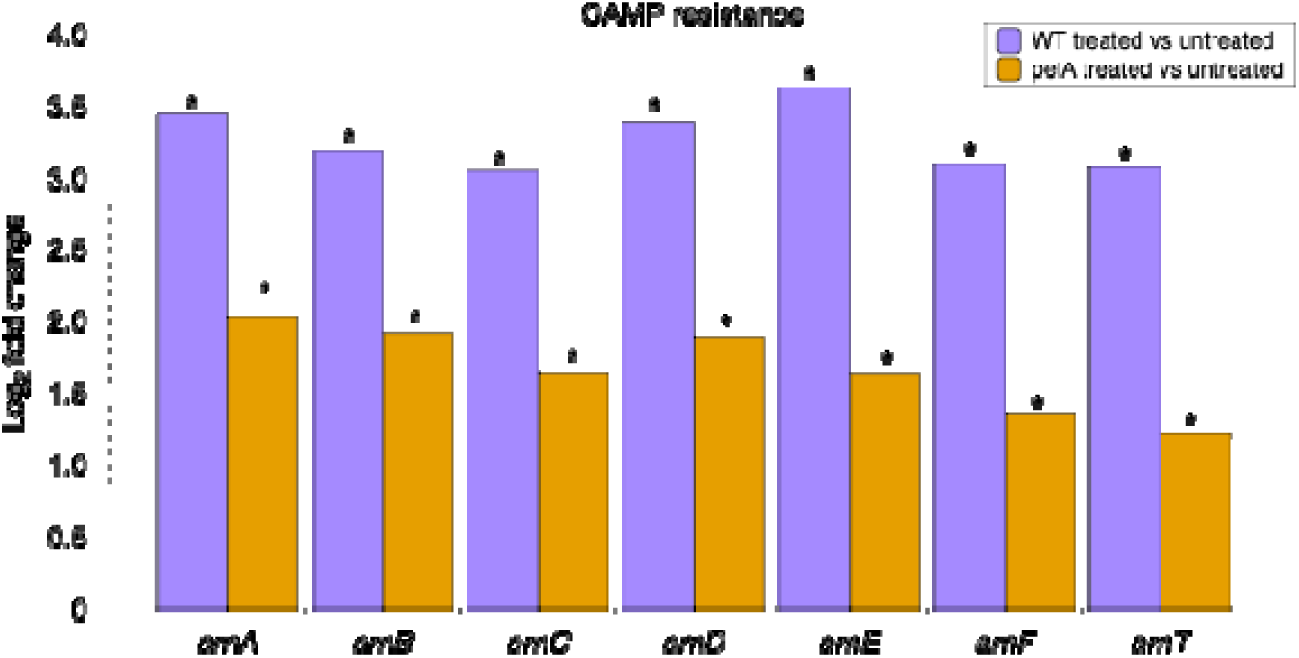
Log_2_ fold change (LFC) in expression of genes of interest related to CAMP resistance by *P. aeruginosa* PA14 WT or *pelA^-^* in the presence vs. absence of colistin. CARD analysis of genes associated with antibiotic resistance from pseudomonas.com. Gene expression analysis was performed between PA14 WT and *pelA^−^* grown in the EVPL, treated and untreated conditions. The dotted line is an LFC of 1.5. The asterisk * denotes an LFC significantly different from zero where the p_adj_ ≤0.05. Gene expression was considered significantly changed when |LFC | was ≥ 1.5 and p_adj_ was ≤0.05. P value corrected for multiple testing with Benjamini and Hochberg method.

## 5 Discussion

It is well established that growth environment affects the antibiotic resistance phenotype of bacteria (Ersoy et al., 2017). Biofilm formation in a chronic infection has been linked with high tolerance to antibiotics and biofilm-deficient transposon mutants, which lack the ability to form a mature, structured biofilm, have been shown to have compromised antibiotic tolerance in *in vitro* biofilm platforms (Colvin et al., 2011). However, we were surprised to find that growth in the EVPL model resulted in a *pelA* transposon mutant exhibiting WT levels of tolerance to colistin and meropenem. Although our results are consistent with two other studies which found that loss of matrix production did not impact antibiotic tolerance if bacterial motility was restricted (Goltermann and Tolker-Nielsen, 2017, Staudinger et al., 2014), we also hypothesised that WT and matrix-deficient mutants might show different responses to the EVPL environment, or to the presence of sub-inhibitory antibiotics, that allowed the mutants to compensate for their lack of matrix. Increases in expression of genes associated with antibiotic resistance were seen in the EVPL vs. SCFM *in vitro*. Similar increases were found in both WT and *pelA^-^*. However, there were only a small number of significant differences in how the two genotypes responded to sub-bactericidal colistin in the EVPL model.

Sub-inhibitory colistin led to changes in expression of genes encoding the MexEF-OprN efflux pump. This set of genes was significantly upregulated by *pelA^−^*in response to treatment but was downregulated by the WT. Expression of *mexEF-OprN* is positively regulated through an activator, MexT (Llanes et al., 2011). Expression of *mexT* was not significantly affected by colistin treatment in either genotype. Thus, another upstream factor likely regulates expression of the pump in our experiment. Upregulation of *mexEF-oprN* is often linked with reduced expression of *oprD*, which encodes an outer membrane porin whose downregulation is associated with resistance to carbapenems and colistin (Lorusso et al., 2022), due to MexT also repressing expression of *oprD*. Consistent with the lack of differential *mexT* expression between WT and biofilm mutants, we did not find significant differential expression of *oprD* in response to colistin between WT and *pelA^−^* mutants in our data set. Although MexEF-OprN itself is not directly associated with antibiotic resistance (Llanes et al., 2011), it is possible that this efflux pump increases tolerance to colistin. It is not usually needed and as such is not normally upregulated – perhaps because, in the WT strain at least, it may decrease fitness of the cells whilst in an anaerobic environment (Olivares et al., 2014). Therefore, it is possible that the loss of pel formation may reduce the costs of the pump, as lack of a matrix results in less oxygen restriction to the cells, removal of the metabolic costs of producing pel, and/or necessitates the increased use of this efflux pump despite a fitness cost. This hypothesis should be explored in future work. It is important to note that the function of efflux pumps is not solely for pumping out antibiotics, but they are also important for the regulation of biofilm formation. Efflux pumps such as MexGHI-OpmD are responsible for transporting quorum sensing molecules including phenazine, which is used for the regulation of biofilm formation (Sakhtah et al., 2016).

## 6 Conclusions and future work

Consistent with our previous study of WT PA14 in the EVPL model, WT biofilms in this study upregulated genes associated with antibiotic efflux and CAMP resistance in response to the EVPL environment, and in response to sub- bactericidal colistin treatment in this model. But unexpectedly, given our previous work showing the resistance of the PA14 biofilm matrix to colistin penetration in EVPL, we found that a lack of polysaccharide in the biofilm does not automatically lead to a reduction in antibiotic resistance in a model where biofilms typically form. When treated with sub-inhibitory colistin concentrations, differences in the types of resistance genes upregulated in biofilm mutants, compared with the WT, may allow them to achieve high levels of resistance.

Overall, we saw similar trends in significant DEGs for the WT PA14 in EVPL vs. *in vitro* as were reported in our previous paper (Harrington et al., 2022). There was a small subset of DEGs that showed the same trend in both studies but only reached statistical or biological significance in one. The expense of RNA sequencing necessarily constrained the size of experiments possible, which reduces statistical power and can lead to between-experiment variation. For instance, we previously reported biologically significant upregulation of genes associated with cationic antimicrobial peptide (CAMP) resistance in PA14 when grown in EVPL vs. SCFM *in vitro*. In the present study, we found statistically significant upregulation of CAMP-resistance-associated loci by both WT and mutant PA14 in the EVPL vs. in SCFM *in vitro,* but not all loci reached the biological significance threshold of a 1.5 LFC. Having demonstrated the tractability of extracting RNA of sufficient quality for this analysis from our EVPL model, and having shown interesting differences in the transcriptome of *P. aeruginosa* grown in the EVPL from SCFM alone, and between genotypes, we hope our results make a solid case for the utility of further work which will enable us to build a larger database of transcriptomes and facilitate high-powered analyses to address a broader range of research questions.

Like most transcriptomics studies, we have sequenced entire populations of cells and reported the average transcriptome for each population; it is entirely possible that subsets of bacteria within each population specialise in expressing different resistance determinants, or that sub-populations with differing levels of metabolic activity achieve intrinsic resistance to different subsets of antibiotics, but these patterns are invisible in the absence of single-cell or spatial transcriptomic approaches (Lyng and Kovács, 2022, Pamp et al., 2008b). Future work using these techniques would provide better understanding of the biofilm population dynamics and their association with antibiotic tolerance.

In conclusion, our results clearly show that despite the physical protection a biofilm matrix can afford from antibiotic attack, biofilm-deficient mutants can still tolerate exceptionally high concentrations of antibiotics in a host-mimicking environment. Further research is needed to tease apart the relative contributions of biofilm matrix and cellular determinants of antibiotic tolerance and resistance to the hyper-resistance phenotypes observed *in vivo*, where biofilm matrix may also be essential for persistence in the presence of a live immune response and cough reflex. Our work suggests that the role of different efflux pumps in determining antibiotic tolerance in wild-type and biofilm-deficient genetic backgrounds is a prime candidate for exploration.

## 7 Acknowledgements

This work was funded by the BBSRC via the Midlands Integrative Biosciences Training Partnership (grant number BB/M01116X/1, PhD studentship awarded to JLL) and by the MRC (grant number MR/R001898/1, New Investigator Research Grant awarded to FH). The CryoSEM performed at the nmRC using the Zeis Crossbeam 550 was supported by the EPSRC (grant number EP/S021434/1). The authors would also like to acknowledge the help of the Media Preparation Facility in the School of Life Sciences at the University of Warwick, with special thanks to Cerith Harries and Charlotte Curtis. We thank four anonymous reviewers for helpful feedback on an early version of this manuscript.

## 8 Author contributions

Conceptualisation: JL, FH. Methodology: FH, NH, SEB, CP. Investigation: JL, DW, RGM, SEB, CP. Formal Analysis: JL. Writing - original draft: JL. Writing - review editing: JL, NH, RGM, DW, CP, SB, FH. Supervision: FH.

