## Supplementary Figures & Tables for "Biofilm-deficient mutants of *Pseudomonas aeruginosa* have wild-type levels of antibiotic tolerance in a model of cystic fibrosis lung infection"

**Gene expression changes to *Pseudomonas aeruginosa* in different environments**

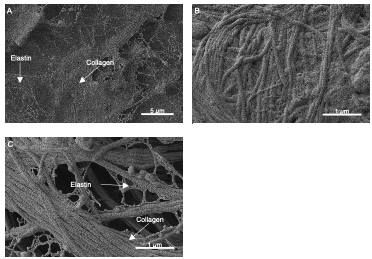

**Figure S1: Cryogenic scanning electron microscopy images of sections of uninfected bronchi tissue extracted from a porcine lung.** Tissue pieces were dissected and washed, then left uninfected surrounded by SCFM for 48 hours in 37 *^◦^*C incubator. Arrows show collagen fibrils with a characteristic banded appearance, and elastin fibres with a characteristic “twisted rope” structure. Images obtained using a Zeiss Crossbeam 550 SEM equipped with a Quorum 3010 Cryo-system performed at 2 kV. Scanning electron microscopy images obtained using a Zeiss Gemini electron microscope with an InLens detector, at 1kV. For a comparative reference image of mammalian collagen fibres, see image 3735 in the National Institute of General Medical Sciences’ online image gallery (<https://nigms.nih.gov/image-gallery/3735>). For comparative reference images of the collagen and elastin network in mammalian lung, see Figure 6 in Wagner et al. (2015) *The Anatomical Record* 298:1960-8; doi: 10.1002/ar.23259.

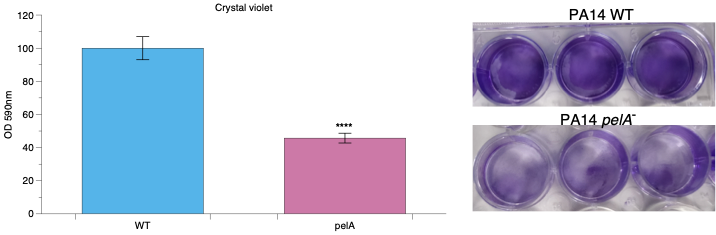

**Figure S2:** **Crystal violet assay staining biofilm mass produced by *P.aeruginosa* PA14 WT and *pelA****^−^*. Left: Bar chart of crystal violet assay showing biofilm mass of *P.aeruginosa* PA14 WT and *pelA^−^*. Right: Image of biofilm stained mass in WT and *pelA^−^* genotypes. PA14 WT and *pelA^−^* were grown in SCFM for 48 hours to form a mature biofilm. Spent media was removed, leaving the biofilm formed on the plate well bottom, crystal violet was added and measured to see overall biofilm mass produced. OD was read at 590nm. Error bars show standard deviation with 6 repeats measured per genotype. The asterisk * denotes a statistically significant result. Two-sample t test was performed (P= *<*0.0001, t=17.56, df-10)

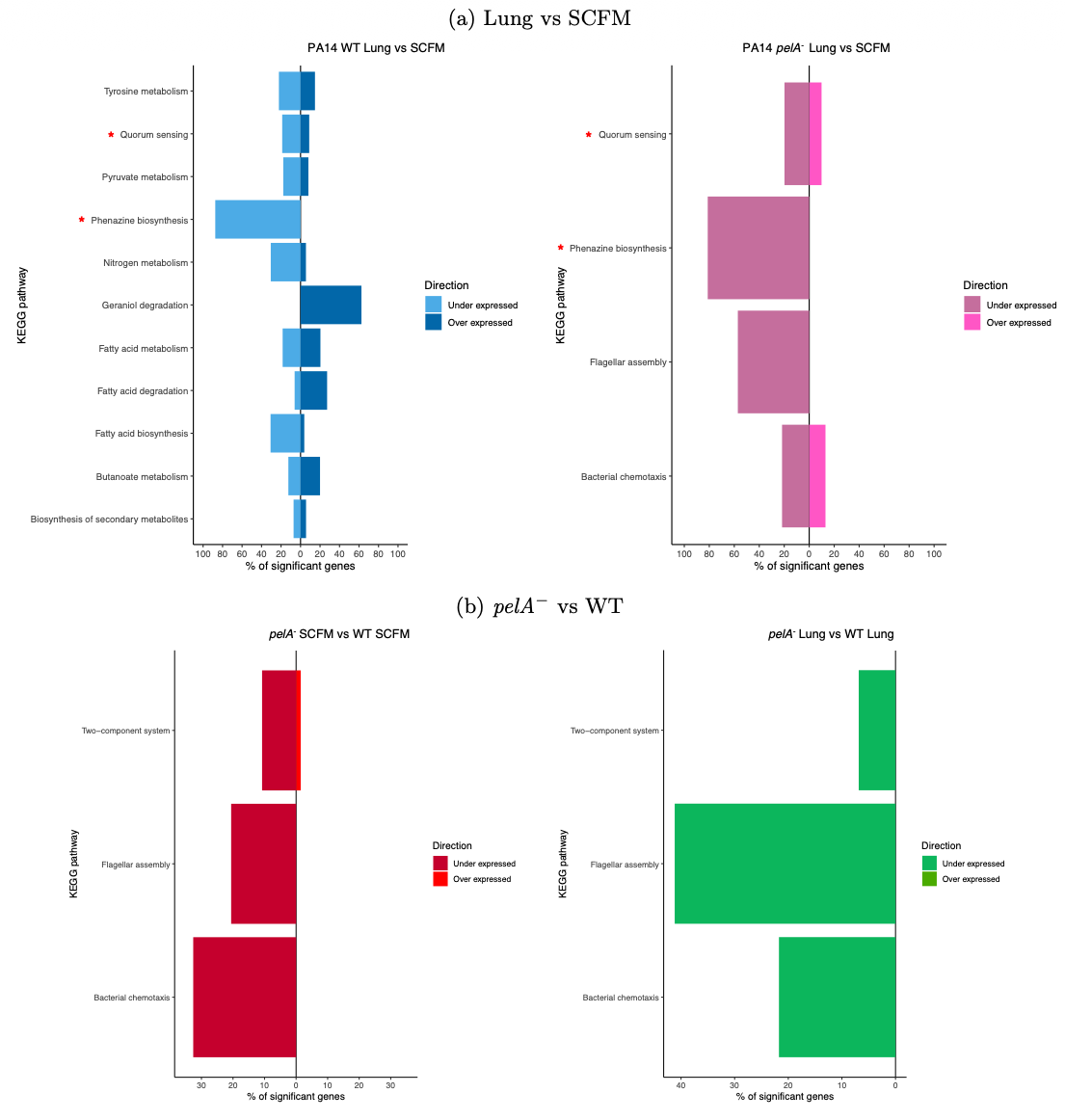

**Figure S3: KEGG analysis of enriched pathways in (a) *P. aeruginosa* PA14 WT or**

***pelA^-^* grown in the EVPL vs. SCFM and (b) *pelA^-^* vs WT in the EVPL or SCFM.** The

bar charts show the percentage of genes that are significantly up- or downregulated in each significantly-enriched pathway. Genes that are upregulated are shown right of the zero line and downregulated genes to the left. Genes were classed as significantly differentially expressed if they had a |Log2 FoldChange| ≥ 1.5 and a p_adj_ < 0.05. The red asterisk show that the pathways are enriched in both genotypes. P-values were corrected for multiple testing with the Benjamini and Hochberg method.

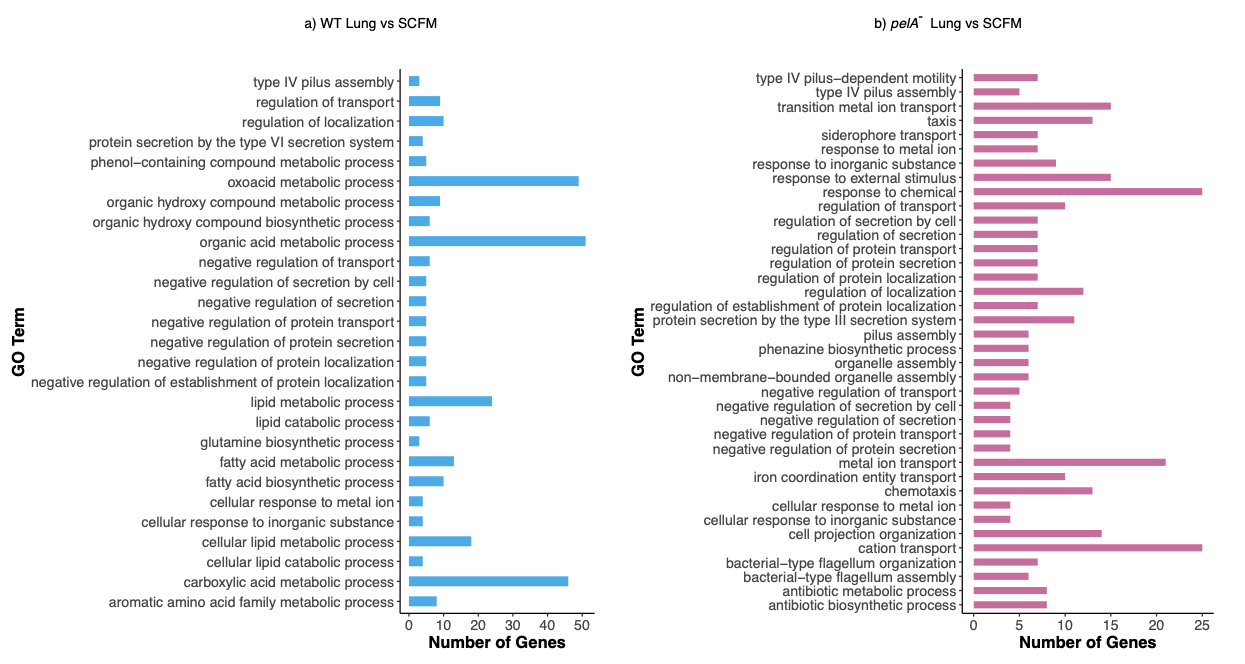

**Figure S4:** **Gene ontology (GO) enrichment analysis of (a) *P. aeruginosa* PA14 WT or (b) *pelA^-^* grown in the EVPL vs. SCFM.** The bar graphs show the significantly enriched GO terms (Fisher’s exact test, p < 0.05) with number of DEGs found for each term. Genes were classed as significantly differentially expressed if they had a |Log2 FoldChange| ≥ 1.5 and a p_adj_ < 0.05. P-values were corrected for multiple testing with the Benjamini and Hochberg method.

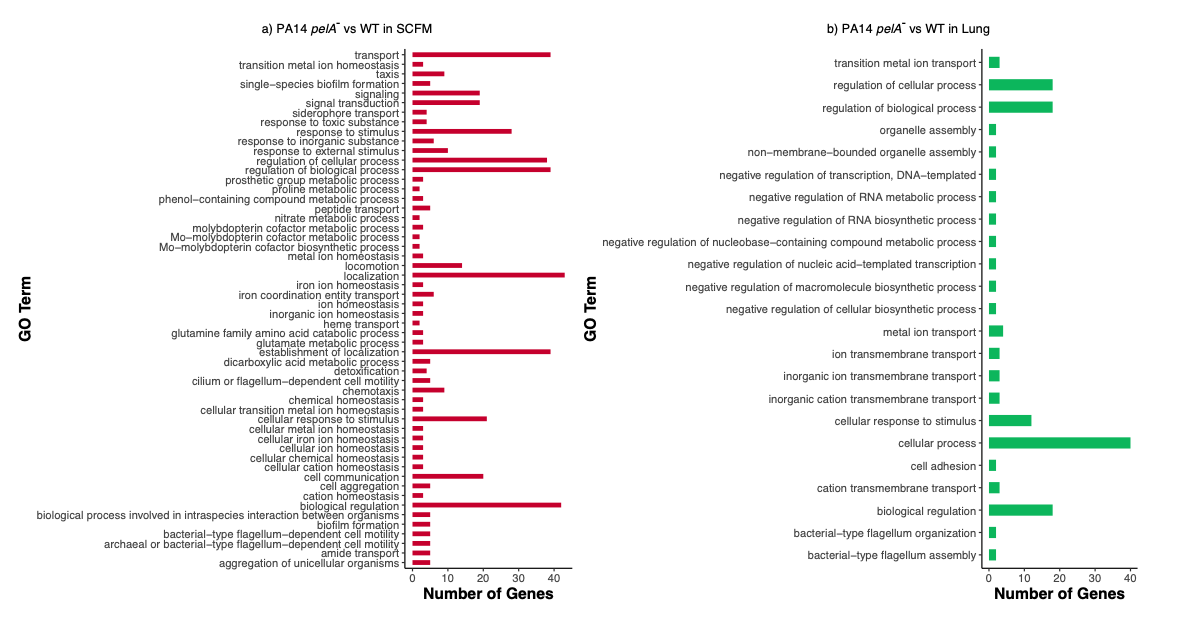

**Figure S5:** **Gene ontology (GO) enrichment analysis of *P. aeruginosa* PA14 WT vs. *pelA^-^* grown in (a) SCFM or (b) EVPL.** The bar graphs show the significantly enriched GO terms (Fisher’s exact test, p < 0.05) with number of DEGs found for each term. Genes were classed as significantly differentially expressed if they had a |Log2 FoldChange| ≥ 1.5 and a p_adj_ < 0.05. P-values were corrected for multiple testing with the Benjamini and Hochberg method.

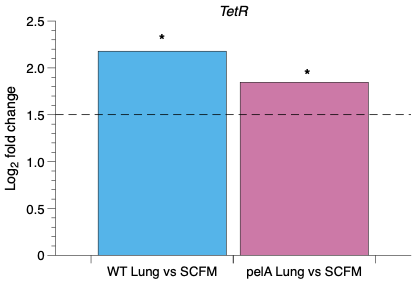

**Figure S6: Log_2_ Fold Change (LFC) of the *TetR* gene in *P. aeruginosa* PA14 WT and *pelA^−^* when comparing growth in the *ex vivo* pig lung model (EVPL) and synthetic cystic fibrosis sputum media (SCFM) *in vitro*.** Black dotted line denotes LFC of 1.5. Gene expression was considered significantly different when *|*LFC*|* was *≥* 1.5*|* and p_adj_ was *≤*0.05. P value corrected for multiple testing with Benjamini and Hochberg method.

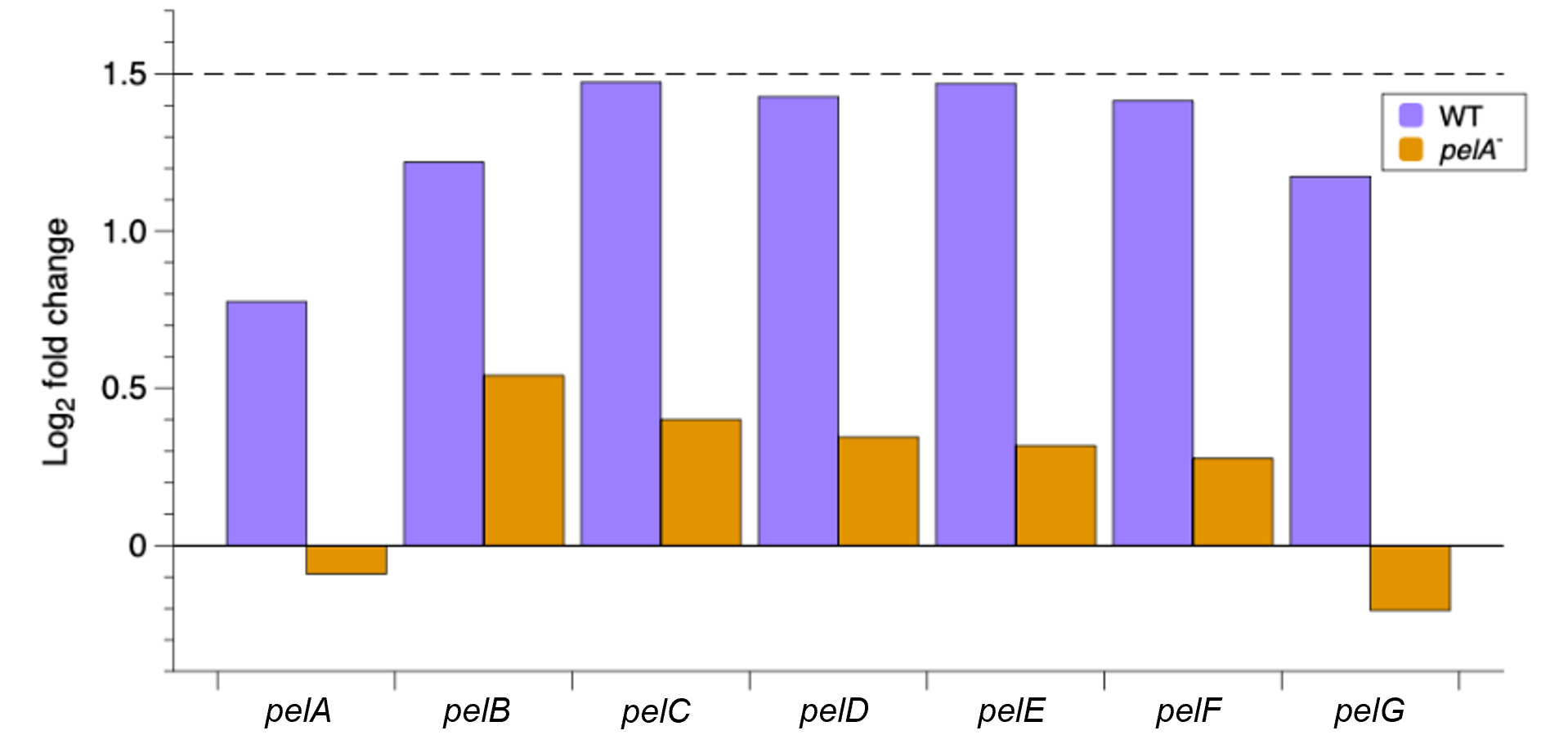

**Figure S7: Log_2_ fold change (LFC) of genes in the *pel* operon in *P. aeruginosa* PA14 WT and *pelA^−^* when comparing growth in the EVPL in the presence vs. absence of sub-bactericidal colistin**. Black dotted line denotes LFC of 1.5. Gene expression was considered significantly different when *|*LFC*|* was *≥* 1.5 and p_adj_ was *≤*0.05. P value corrected for multiple testing with Benjamini and Hochberg method. No loci were significantly differentially expressed by either genotype.

| GeneID | Gene name | WT  EVPL vs SCFM | *pelA^−^*  EVPL vs SCFM | EVPL  *pelA^−^* vs WT | SCFM  *pelA^−^* vs WT |
| --- | --- | --- | --- | --- | --- |
| PA14 18380 | *algA* | 1.41 | 0.17 | -1.07 | 0.17 |
| PA14 70270 | *algC* | -0.75 | -0.44 | 0.04 | -0.27 |
| PA14 18580 | algD | **2.07*** | 1.10 | -0.99 | -0.02 |
| PA14 18510 | *algE* | 0.88 | 0.53 | -0.26 | 0.08 |
| PA14 18410 | *algF* | 1.16 | -0.27 | -0.87 | 0.55 |
| PA14 18500 | *algG* | 0.27 | -0.30 | -0.26 | 0.31 |
| PA14 18450 | *algI* | 0.92 | 0.54 | -0.48 | -0.10 |
| PA14 18430 | *algJ* | 1.05 | 0.41 | -0.52 | 0.12 |
| PA14 18520 | *algK* | 0.43 | 0.84 | -0.06 | -0.47 |
| PA14 18470 | *algL* | 0.90 | 0.26 | -0.54 | 0.10 |
| PA14 18565 | *alg8* | 1.17 | 0.86 | -0.37 | -0.06 |
| PA14 18550 | *alg44* | 0.79 | 0.74 | -0.40 | -0.35 |
| PA14 18480 | *algX* | 1.02 | 0.85 | -0.33 | -0.16 |

**Table S1.** **Log_2_ FoldChange (LFC) of genes included in the alginate biosynthesis pathway in *P. aeruginosa* PA14 WT and *pelA****^−^* **in the EVPL vs. in SCFM.** Significantly differentially expressed genes are denoted with an asterisk * and are classed as having |LFC| of *≥* 1.5 and P < 0.05. P values were corrected for multiple testing with the Benjamini and Hochberg method.

| GeneID | Gene name | WT  treated vs. untreated | *pelA^-^*  treated vs. untreated |
| --- | --- | --- | --- |
| PA14 18380 | *algA* | -0.22 | 0.10 |
| PA14 70270 | *algC* | 0.34 | -0.01 |
| PA14 18580 | *algD* | 0.12 | 0.02 |
| PA14 18510 | *algE* | -0.17 | 0.34 |
| PA14 18410 | *algF* | -0.11 | -0.07 |
| PA14 18500 | *algG* | -0.24 | -0.04 |
| PA14 18450 | *algI* | -0.20 | 0.01 |
| PA14 18430 | *algJ* | -0.16 | 0.19 |
| PA14 18520 | *algK* | 0.34 | 0.43 |
| PA14 18470 | *algL* | 0.08 | -0.03 |
| PA14 18565 | *alg8* | -0.07 | 0.18 |
| PA14 18550 | *alg44* | 0.17 | 0.49 |
| PA14 18480 | *algX* | 0.07 | 0.24 |

**Table S2:** **Log**_2_ **fold change (LFC) in expression of genes involved in alginate biosynthesis in *P. aeruginosa* PA14 WT and *pelA****^−^* **in the presence vs. absence of colistin**. Significantly differentially expressed genes are classed as having |LFC| of *≥*1.5 and P < 0.05. P values were corrected for multiple testing with the Benjamini and Hochberg method. No loci were significantly differentially expressed.

| Gene ID | Gene name | WT treated vs untreated | *pelA^-^* treated vs untreated |
| --- | --- | --- | --- |
| PA14_18300 | Probable nucleotide sugar dehydrogenase | 2.33* | 0.73 |
| PA14_18340 | *arnD* | 3.38* | 1.89* |
| PA14_18350 | *arnA* | 3.44* | 2.02* |
| PA14_18360 | *arnC* | 3.04* | 1.63* |
| PA14_18370 | *arnB* | 3.18* | 1.91* |

**Table S3: Log_2_ FoldChange (LFC) of genes included in the amino sugar and nucleotide sugar metabolism GO term in *P. aeruginosa* PA14 WT and *pelA****^−^* **in the presence vs. absence of colistin** Significantly differentially expressed genes are denoted with an asterisk * and are classed as having |LFC| of *≥* 1.5 and P < 0.05. P values were corrected for multiple testing with the Benjamini and Hochberg method.
